# Drivers of host-infectious agent community associations in seabirds from sub-Antarctic oceanic islands

**DOI:** 10.64898/2026.02.23.706941

**Authors:** Tristan Bralet, Mathilde Lejeune, Jérémy Tornos, Augustin Clessin, Clémence Galon, Florian Berland, Alexandre Mokorel-Pouye, Amandine Gamble, Sara Moutailler, Rachid Aaziz, Karine Laroucau, Thierry Boulinier

## Abstract

Understanding the patterns and drivers of infectious disease dynamics at different levels of life organization is a fundamental disease ecology challenge. In this study, we performed a multi-infectious agents (IA) screening of 1,983 individuals belonging to 18 seabird species, sampled between 2017 and 2022 across five sub-Antarctic islands in the Southern Indian and Atlantic Oceans. Our aim was to identify the drivers of IA community in these seabirds by considering intrinsic and extrinsic factors on three dimensions: distance between seabird metapopulations and within-island sites, macro-community (host functional traits based on feeding and breeding features) and intra-host IA community composition. Samples were screened for 24 DNA-based IAs using a high-throughput real-time PCR method, followed by subsequent hierarchical modelling of species communities. *Campylobacter lari*, *Escherichia coli, Pasteurella multocida, Chlamydiaceae* and *Mycobacterium* spp. were found to be present on all islands in some, but not all, species. However, only *E. coli* was detected in all species. According to our modelling results, the influence of year and within-island site had limited influence on IA presence/absence, accounting respectively for an average of 1.0% and 1.8% of the variation in IA community. Including host species as a response variable produced a better model fit than including host species’ functional traits, highlighting the difficulty in simplifying systems by focusing on high-level functional categories. Model outputs show how each species influences IA distribution depending on IA traits, but no clear trend has emerged. Interestingly, burrowing species were less frequently infected with directly transmitted IAs, probably due to the low frequency of transmission events associated with their breeding features and limited social contacts. Our findings pave the way for further multi-dimension studies that would combine complementary approaches to disentangle the complex processes at play in the dynamics of hosts and IAs interactions.

## INTRODUCTION

Understanding the drivers of infectious processes and disease emergence is critical in the current context of global change for addressing conservation issues, global health concerns and economic values (Dobson *et al*., 2020; Gupta *et al*., 2020). This requires answering key eco-epidemiological questions, such as which how infectious agents (IAs) circulate, which one could emerge, and which populations could be threatened (Daszak *et al*., 2000). A key challenge in disease ecology is understanding the importance of biotic and abiotic drivers underlying these processes at all levels of life organization, from meta-ecosystems to individuals (Tompkins *et al*., 2011; Cohen *et al*., 2016; Stephens *et al*., 2016; Elderd *et al*., 2022).

At large scale, from a macroecology perspective, exploring the factors that mostly influence IA diversity and distribution is of major interest (Cohen *et al*., 2016; Talmon *et al*., 2025). These factors include abiotic conditions, but also large-scale host movements and anthropisation. Further, it is important to understand which scale (regional, global,…) can be used to identify general trends or variability in IA distribution and communities (Levin, 1992). On another dimension, the host community (“macro-community”), the challenge is to understand how IAs are shared between host species (Locke *et al*., 2013; Fecchio *et al*., 2017), what causes differences in exposure or susceptibility between host species (Sweeny & Albery, 2022), and which species act as reservoirs (Worsley-Tonks *et al*., 2020) or spatial spreaders of each IA, depending on its transmission and life cycle (Gamble *et al*., 2020; Manlove *et al*., 2022).

Finally, at the host level, each individual may harbor a specific assemblage of IAs (“infra-community”). The factors that determine these combinations, as well as the direct or indirect interactions between IAs, are of particular interest (Sáez-Ventura *et al*., 2022), as are the effects that IA communities have on their host’s health.

Only a few studies have so far incorporated multiple dimensions within the same analysis (Moss *et al*., 2020) investigating the influence of different biological dimensions on the structure of IA communities. Furthermore, a multi-dimension approach requires an appropriate framework and adapted analytical methods (Tompkins *et al*., 2011; Elderd *et al*., 2022). However, such ecological studies frequently encounter complex multi-host systems where interactions often exceed the scope of conventional multi-site, multi-IAs investigations, hindering the establishment of strong causal inferences (Caron *et al*., 2015). To address this, disease ecologists often simplify systems by focusing on high-level functional rather than species-level taxonomic diversity (Hooper *et al*., 2002; Jax, 2005). Defining functional groups requires the initial characterization of host functional traits (see Caron *et al*., (2012) for the definition of functional groups in a study on Influenza virus circulation in birds), after which statistical frameworks can be employed to identify traits of particular epidemiological relevance (Plowright *et al*., 2008; Caron *et al*., 2015). For example, such a trait-based approach can help to identify candidate species that may act as spreaders (Tornos *et al*., in prep) or reservoirs (Worsley-Tonks *et al*., 2020).

Due to their simple biotas and clear spatial structure, insular systems are ideal models for studying ecological processes (Kueffer *et al*., 2014; Fountain-Jones *et al*., 2024). Seabird are of particular interest in the context of disease ecology due to their unique characteristics (Bralet *et al*., in prep). For example, their colonial breeding exposes them to local transmission of IAs, while their long-range movements could facilitate IA spread between islands. In light of the increasing awareness of the impact of epizootics on seabird populations, which are already under the threat of global change and human activities, it is essential to investigate infectious processes for conservation purposes (Dias *et al*., 2019). This has been the case with avian cholera (e.g., in common eiders (*Somateria molissima)* in the Arctic and in Indian yellow-nosed albatrosses (*Thalassarche carteri)* on Amsterdam Island (Descamps, *et al*., 2011; Jaeger *et al*., 2018), Southern Indian Ocean), but also more recently as highlighted in the panzootic of high pathogenicity avian influenza (Knief *et al*., 2024; Lane *et al*., 2024).

An interesting feature of subantarctic insular systems is that they share relatively simple and comparable species assemblages in terms of both phylogeny and ecological traits, even when situated at great distances from one another (Delord *et al*., 2014). This is likely due to their similar abiotic conditions, such as climate and geology (Moon *et al*., 2017), as well as their common history and host speciation processes (Inchausti & Weimerskirch, 2002; Clucas *et al*., 2018; Pertierra *et al*., 2020).

Host communities can therefore be compared at various spatial scales, both within and between oceanic basins, with for instance a common set of scavengers at colonies and densely breeding species above or underground. Groups of islands in a given area can be considered as spatial ‘replicates’ in order to answer certain questions. However, issues relating to statistical non-independence must be taken into account (Selmi & Boulinier, 2001), particularly when considering spatial scales at which seabird movements could be significant (Moon *et al*., 2017).

In this context, we aimed to explore the IA communities of seabirds on sub-Antarctic islands to identify potential drivers of infectious status of hosts by using a trait-based approach. These drivers included large-spatial-scale extrinsic factors (e.g., the distance between metapopulations or populations), host-community-dimension factors (e.g., the ecological and functional features of hosts, such as their trophic position, breeding density and breeding habits) and infra-community-dimension factors (e.g., the mode of transmission and co-infections). To this end, we screened 1,983 seabirds from five subantarctic territories at various distances for 24 IAs. We used an adapted high-throughput real-time PCR method alongside a hierarchical modelling of species communities (HMSC) framework to test the following hypotheses:

i. <u>at the Southern Ocean scale</u>, we expected the qualitative composition of the IA community (e.g. the presence of each IA) to be similar between islands with comparable climatic conditions and seabird communities;
ii. <u>at the island scale</u>, we did not expect to observe any quantitative differences in IA community composition between sites within an island, particularly given the high level of seabird mobility between colonies;
iii. <u>at the host-community dimension</u>, we expected greater IA diversity and prevalence in terrestrial predators and scavengers than in other taxa, as they are supposedly repeatedly exposed to IAs while preying on other species. Furthermore, we anticipated greater IA diversity and prevalence in ground-breeding mesopredators than in burrow-nesting species due to their reduced exposure and social interactions through burrow-nesting behavior;
iv. <u>at the infracommunity dimension</u>, we expected a higher proportion of directly transmitted IAs to be present in the IA communities of predators and scavengers, as well as in high-density ground-breeding species, due to facilitated transmission. Conversely, we expected to see more environmentally-transmitted IAs in burrow-

nesting species. We also expected a positive co-occurrence between IAs that share the same transmission mode.

## MATERIAL AND METHODS

### 1. Samples collection

The data were collected through field sampling campaigns on four subantarctic islands in the South Indian Ocean and one in the South Atlantic Ocean : Possession Island (46°25’S, 51°45’E) and Cochons Island (49°28’S, 70°2’E) in the Crozet Archipelago, the Kerguelen Islands (49°15’S, 69°10′E), Amsterdam Island (37°49′S, 77°33′E), and New Island (51°43′S, 61°18′W) in the Falkland/Malvinas Islands (POS, COC, KER, AMS and NWI, respectively).

Sampling either took place during a single field campaign (in 2018/19 for NWI and KER, and in 2019/20 for COC), or was conducted over several years to investigate potential temporal variations in epidemiological processes (in 2021/22 and 2022/23 for AMS, and in 2017/18, and 2022/23 for POS). Each sampling period took place during the austral summer.

Sampling was carried out at several sites (colonies) on each island to allow for spatial replication (**Supplementary Material S1&S2**). A total of 1,709 apparently healthy adult seabirds belonging to 18 species were captured by hand (for penguins) or with a noose pole or hook (for most flying birds) before being sampled using a cloacal swab.

The swabs were stored either in Longmire preservation buffer (Longmire *et al*., 1997) or RNA Later (Qiagen), at cool ambient temperatures for up to one week and then at - 20°C.

### 2. Screening for infectious agents

Nucleic acids were extracted from the swabs using the method described by Bralet *et al*., (2025). Searches for IA in each swab were performed using a high-throughput microfluidic real-time PCR (Htrt PCR) adapted from Bralet *et al*., (2025). Briefly, the Htrt PCR uses a chip capable of performing 2,304 of simultaneous single-plex PCR reactions. This enables up to 48 samples to be simultaneously screened for up to 48 IAs.

We adapted the PCR assay by adding and removing PCR systems to search for the following IAs of interest in seabirds: the new genus *Chlamydiifrater* spp., six species of *Enterococcus* spp. (see below), *Escherichia coli* and three Shiga toxin-producing *E. coli* (STEC) pathotypes (see **Supplementary Material S3** for details on PCR systems). We also included a PCR system targeting the avian and mammalian ß-actin gene (ID-VET, Grabels, France) as an extraction control. One negative water control and two positive controls, consisting of a mixture of all positive control DNA divided into two wells, were included per chip. Ct values >30 were considered negative. Overall, this method enabled us to screen 45 samples for 36 bacterial, fungal and protozoal taxa (**Supplementary Material S4**). We made the assumption that there were no differences in conservation features between preservation buffers. For the prevalence calculation and subsequent modelling steps, we also included data from Bralet *et al*., (2025), which screened 274 seabird samples from POS (2021/22) for a slightly different list of targeted IAs.

### 3. Data analysis

Analyses were performed using CRAN R v4.3.3 (R Core Team, 2021).

#### 3.1 Host species grouping

Seabird species were grouped according to their ecological traits (e.g. trophic level, feeding area, breeding density and breeding type), based on the prediction presented above, using principal component analysis (**Supplementary Material S5**). The following functional groups were then identified: (i) terrestrial apex predators and scavengers (skuas, giant petrels and sheathbills); (ii) ground-nesting coastal birds (gulls and shags); (iii) ground-nesting mesopredators (penguins and albatrosses) and (iv) burrow-nesting mesopredators (small petrels).

#### 3.2 Grouping of IA traits

Following a review of the literature (**Supplementary Material S6**), IAs were categorized according to their primary mode of transmission (direct or environmental).

#### 3.3 Prevalence and diversity estimation

The true prevalence and 95% Clopper-Pearson confidence intervals for each IA were estimated using the ‘prevalence’ R package (Devleesschauwer *et al*., 2022) assuming test specificity and sensitivity of 1 and 0.95, respectively (Bralet *et al*., 2025). IA diversity was estimated by calculating specific richness and the Shannon index using the ‘vegan’ package (Oksanen *et al*., 2025). To avoid redundancy, we only considered the results of PCR sets targeting the broadest taxonomic ranks for each IA, acknowledging inequality in taxonomic resolution among taxa. These included the following 24 taxa: *Brucella* spp., *Campylobacter coli*, *Campylobacter jejuni*, *Campylobacter lari*, *Chlamydiaceae*, *Coxiella burnetii*, *Enterococcus casseliflavus, Enterococcus faecalis*, *Enterococcus faecium*, *Enterococcus gallinarum*, *Enterococcus hirae*, *Enterococcus mundtii*, *Erysipelothrix amsterdamensis*, *Erysipelothrix rhusiopathiae*, *Escherichia coli*, *Leptospira* spp., *Mycobacterium* spp., *Pasteurella multocida*, *Salmonella* spp., *Yersinia* spp., *Aspergillus fumigatus*, *Fusarium* spp., Mucorales and *Toxoplasma gondii*.

#### 3.4 Statistical analysis

The factors potentially associated with infection patterns, such as host’s functional group, species and sampling sites, were investigated using a joint species distribution model (JSDM) (Ovaskainen *et al*., 2017). This hierarchical Bayesian framework based on a multivariate generalized linear mixed model with hierarchical layers was used to analyse IAs communities, accounting for fixed and random covariates, as well as IA traits (see Ovaskainen *et al*., (2017) for model formulation). The response variable was the binomial presence-absence of each of the IAs in the hosts. We opted for intrinsic (s*pecies* or *functional group*) and extrinsic (*year*, *island* and *within-island sampling site* (*WIS*)) explanatory covariates. The extrinsic covariates were considered as random effects, as individuals sampled in the same year and site may share similar infectious status. We also incorporated the specific transmission mode of each IA into the trait matrix.

As expected in such multi-scale studies, not all species are sampled at all locations and times due to logistical constraints, hence there is some artefactual dependence between some explanatory variables. Depending on islands or year, only species of certain functional groups could be sampled, and only some host species were sampled at multiple sites within an island. A model based on multi-species data from all islands could not be run because only one species (the brown skua, *Stercorarius antarcticus*) was sampled in all the islands. Moreover, the host species and functional group variables are dependent, as the host species fully determines the functional group.

We therefore adopted a sequential approach, running three series of models for which the predictors were independent (**Table 1**). These corresponded to three spatial scales: (1) “island-scale”, estimating the effects of *year* and *WIS* based on seabird species sampled in at least two POS campaigns (2017, 2021 and 2022); (2) “regional-scale”, estimating the effect of *functional group* on IA communities using data from species shared across POS (2022), COC (2019) and KER (2018) within the same ocean; and (3) “global-scale”, estimating the effect of *functional group* on IA community’s species common to distant islands (KER (2018), AMS (2022) and NWI (2018)) between two oceans. To test the effect of *host species* vs. *functional group* on the IA presence/absence, we ran separate models, each including only one of these two variables as a fixed effect. We then compared the models’ explanatory values and outputs. Since WIS explained only 1.0% of the variance in the island-scale models (**Figure 4**), we addressed the issue of collinearity between site and host species in the regional- and global-scale models by assuming there was no effect of WIS and treated *island* and *year* as random effects (**Table 1**).

**Tableau 1.**
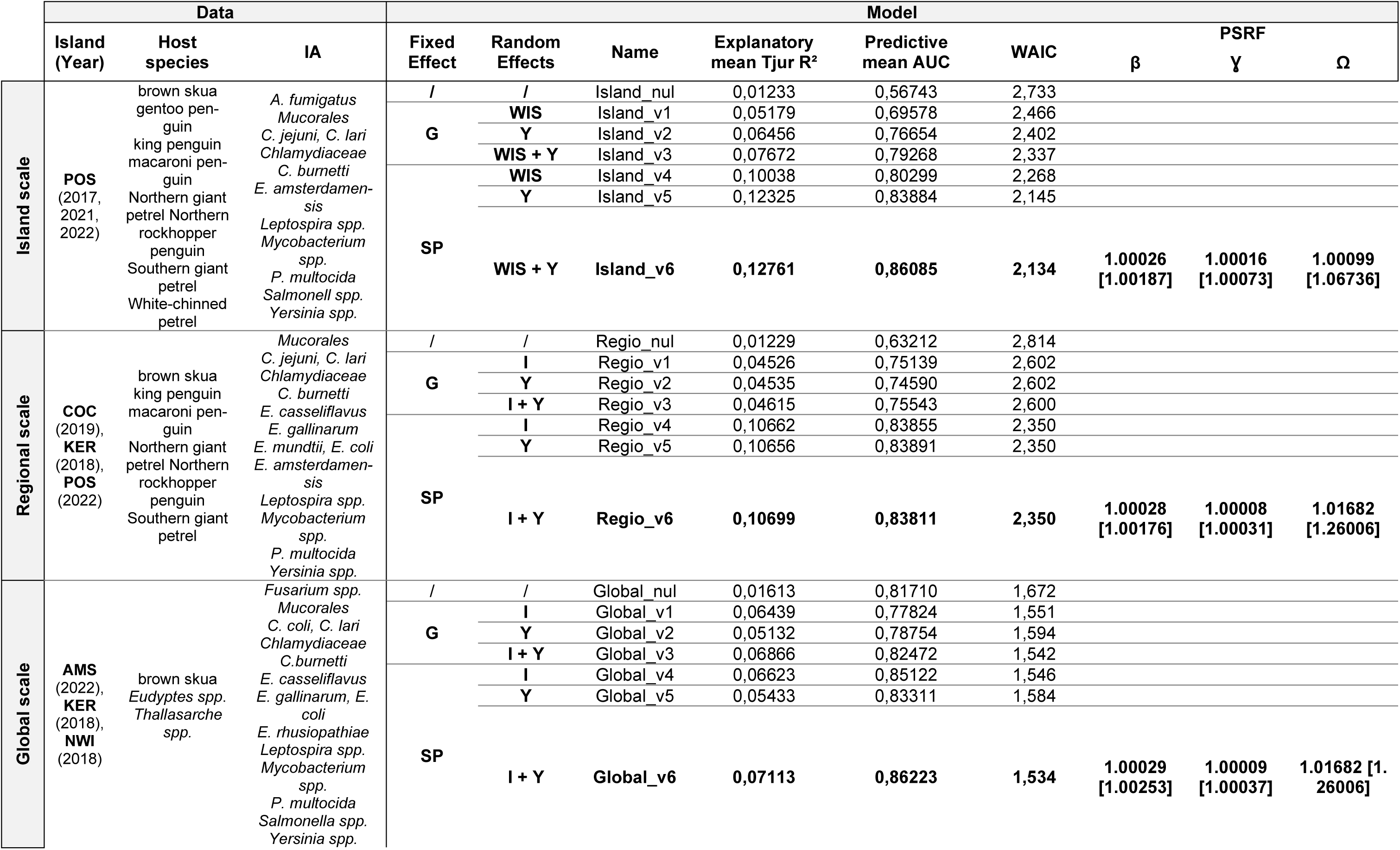
Model characteristics and performance to account for multiple infection patterns. Conditions and data (selected IA, islands, field campaign year and host species) of each of the three series of models (Island-, Regional- or Global-scale) are specified. Fixed effects: G: Ecological group, S: Host species / Random effects: I: Island, WIS: Within-island site, Y: Year of sampling. The final column provides the mean potential scale reduction factors and maximum values (in brackets).

For each series, we compared the most complex model with a simpler one by removing extrinsic variables. Only IAs with at least one occurrence and seabird species sampled at a minimum of two sites were included in each analysis (**Table 1**).

Model outputs include 3 types of coefficients. **β** denotes the response of presence-absence of each of the IAs to host-related variables : each β coefficient associated with a qualitative variable (here, the *host species* or host *functional group*), measures the average displacement, in standard deviations on the normal latent scale, between the considered category and a reference category (“intercept”: king penguin in the island- and regional-scale models, *Eudyptes* spp. for the global-scale model). A positive β value indicates a higher probability of occurrence, and conversely for negative value. **Ɣ** represent the contribution of the IA trait (here, the transmission mode) to this difference. **Ω** describes the residual co-occurrence patterns among AIs at each random effect (*year, WIS, island),* i.e. whether IAs tended to co-occur within years, sites and islands.

The JSDM was fitted to the IA community data using the ‘Hmsc’ R package v. 3.0–14 (Tikhonov *et al*., 2020). The models were fitted using Bayesian inference through Markov chain Monte Carlo (MCMC) method, as described by Berland *et al*., (2025). Default prior distributions were employed, and posterior distributions were obtained using four independent MCMC analyses, each comprising 1,600,000 iterations. The first 200,000 iterations were discarded as burn-in and the chains were thinned at hundred-iteration intervals. Convergence was assessed visually and via potential scale reduction factor with the ‘ggmcmc’ R package v1.5.1.1 (Fernández-i-Marín, (2016).

The quality of the models (explanatory and predictive values) was evaluated using the WAIC (Watanabe, 2010), Tjur’s explanatory R² (Tjur, 2009) and AUC (Pearce & Ferrier, 2000) values. Models with high Tjur’s R² and AUC and low WAIC were selected (Ovaskainen & Abrego, 2020).

The existence of an effects was based on posterior support value (PS), which represents the proportion of posterior distribution that a is strictly positive or negative (Makowski *et al*., 2019; Ovaskainen & Abrego, 2020). For the β and γ parameters, effects were considered non nul when 0 was not within the 95% credible interval. For the Ω parameter, we considered co-occurrence between IAs when the PS was ≥0.95 an “tendency” for PS ≥0.75 (Berland *et al*., 2025).

### 4. Ethical approval

Sampling of live animals in the French Southern and Antarctic Lands was authorised by the French Ministry of Research (APAFIS #31773-2019112519421390 v4) and the TAAF administration (A-2021-55)), following evaluation by Regional Ethics Committee (CREA34), the Polar Environment Committee and the National Council for the Protection of Nature. In the Falkland Islands, the field research was conducted under license no. R28/2018, granted to TBo by the Falkland Islands Government.

## RESULTS

### 1. Infection and co-infection frequencies

Of the 1,709 seabird samples with the new Htrt PCR assay, 62.6% were positive for at least one IA. Overall, 22 IAs were detected, including nine in ≥ 50 samples and three in >10% of all samples (*E. coli*, *Chlamydiaceae* and *Chlamydiifrater* spp.). Five IAs (*C. lari, Chlamydiaceae, E. coli, Mycobacterium* spp.*, P. multocida*) were found on all islands, while 17 occurred on at least two islands and four were restricted to a single island (*C. coli, E. faecium, E. mundtii* and *E. rhusiopathiae)* (**Figures 1 & 2**).

**Figure 1.**
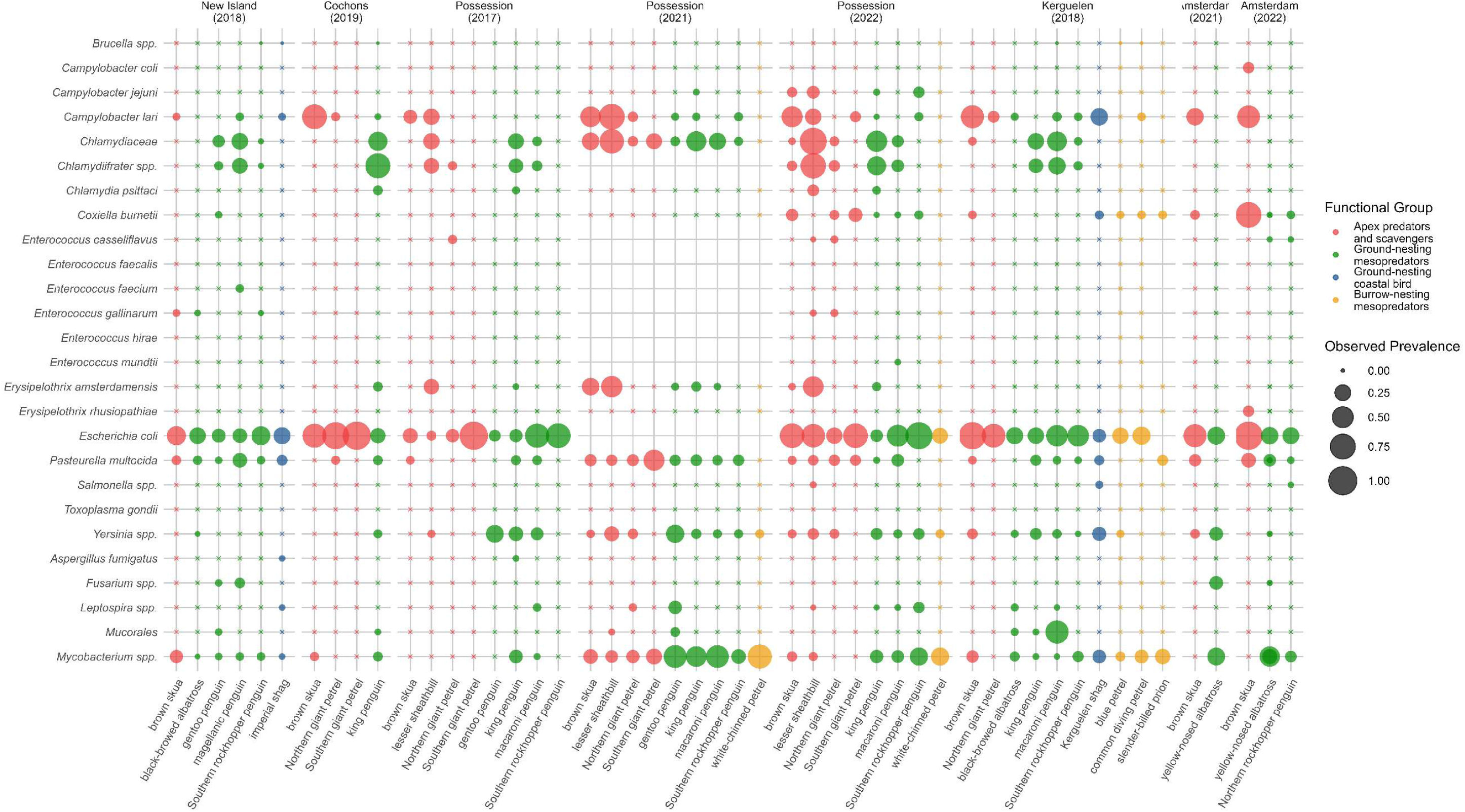
Observed prevalence of infectious agents in seabirds from five different subantarctic territories. Crosses represent zero prevalence.

*E. coli* was the most prevalent IA and was detected in all seabird species and populations; however, none were STEC pathotypes. In contrast, bacteria from the genus *Enterococcus*, thought also to be common in birds’ microbiota, were rarely detected. *P. multocida* was detected at many locations and on many species, notably with relatively high prevalence in the Falkland Islands (reaching 35.1% [9.0–68.2] or 23.9% [8.6–43.8] in some brown skua clubs or penguin colonies) (**Figure 2**).

**Figure 2.**
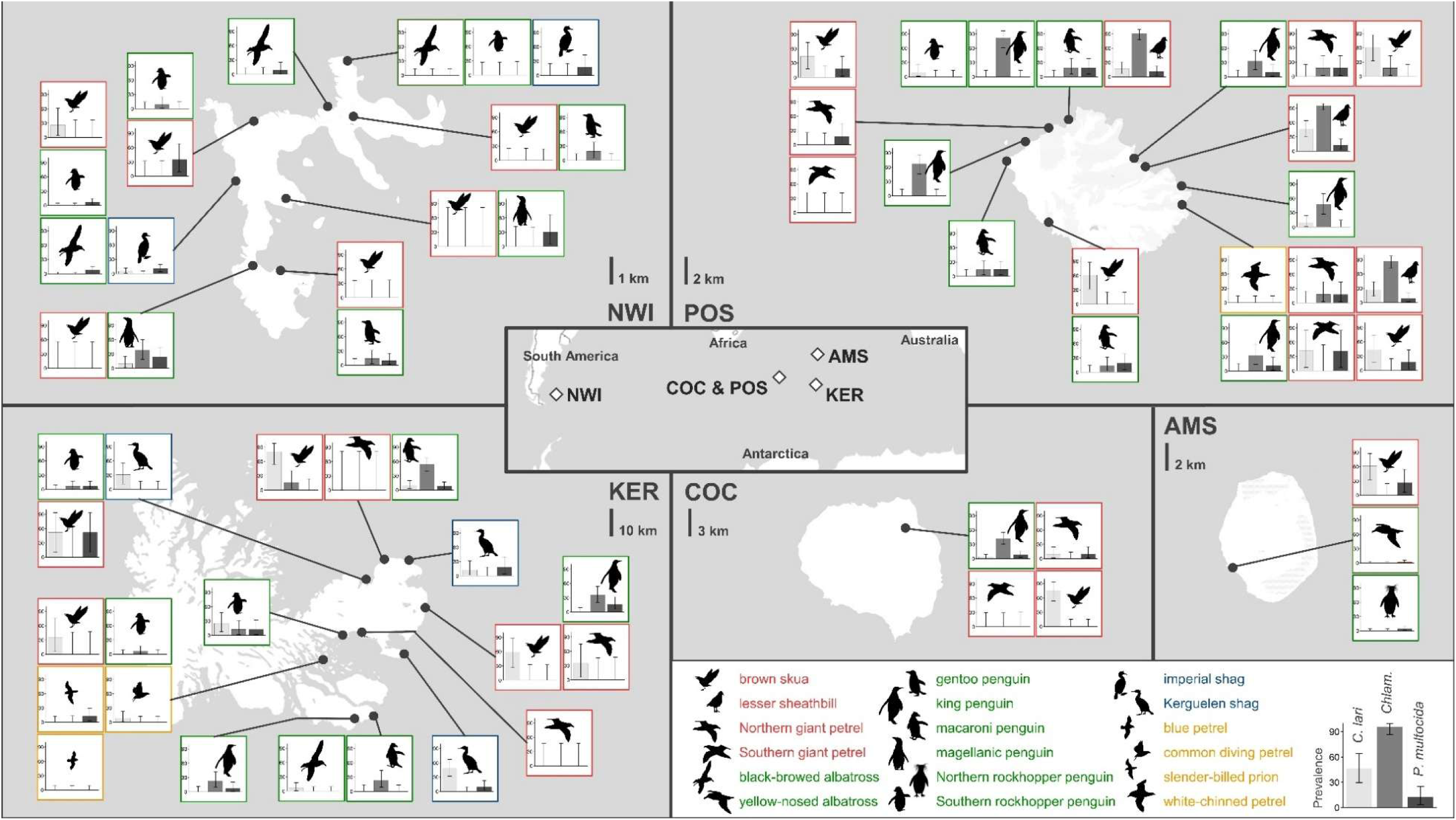
Epidemiological map for five subantarctic territories showing prevalence of IA in seabirds from different ecological groups using Htrt PCR on cloacal swabs. AMS, COC, KER, NWI, POS stands for field campaigns conducted on Amsterdam Island (2022), Cochons Island (2019), Kerguelen Islands (2018), New Island (2018) and Possession Island (2022), respectively. Three IAs of particular interest are represented: *Campylobacter lari* (light grey), *Chlamydiaceae* family (medium grey) and *Pasteurella multocida* (dark grey). See **Supplementary Material S1 & S2** for details on sample sizes and sampling sites.

Most birds carried none of the targeted IAs or only one, and multiple co-infections (up to six IAs) were very rare. Richness ranged from 0 (38.9% of samples; 646 individuals) to 6 (0.2% of samples: two lesser sheathbills (*Chionis minor*) and one macaroni penguin (*Eudyptes chysolophus*)). The median and mean richness were 1 and 0.95, respectively (**Figure 3**).

**Figure 3.**
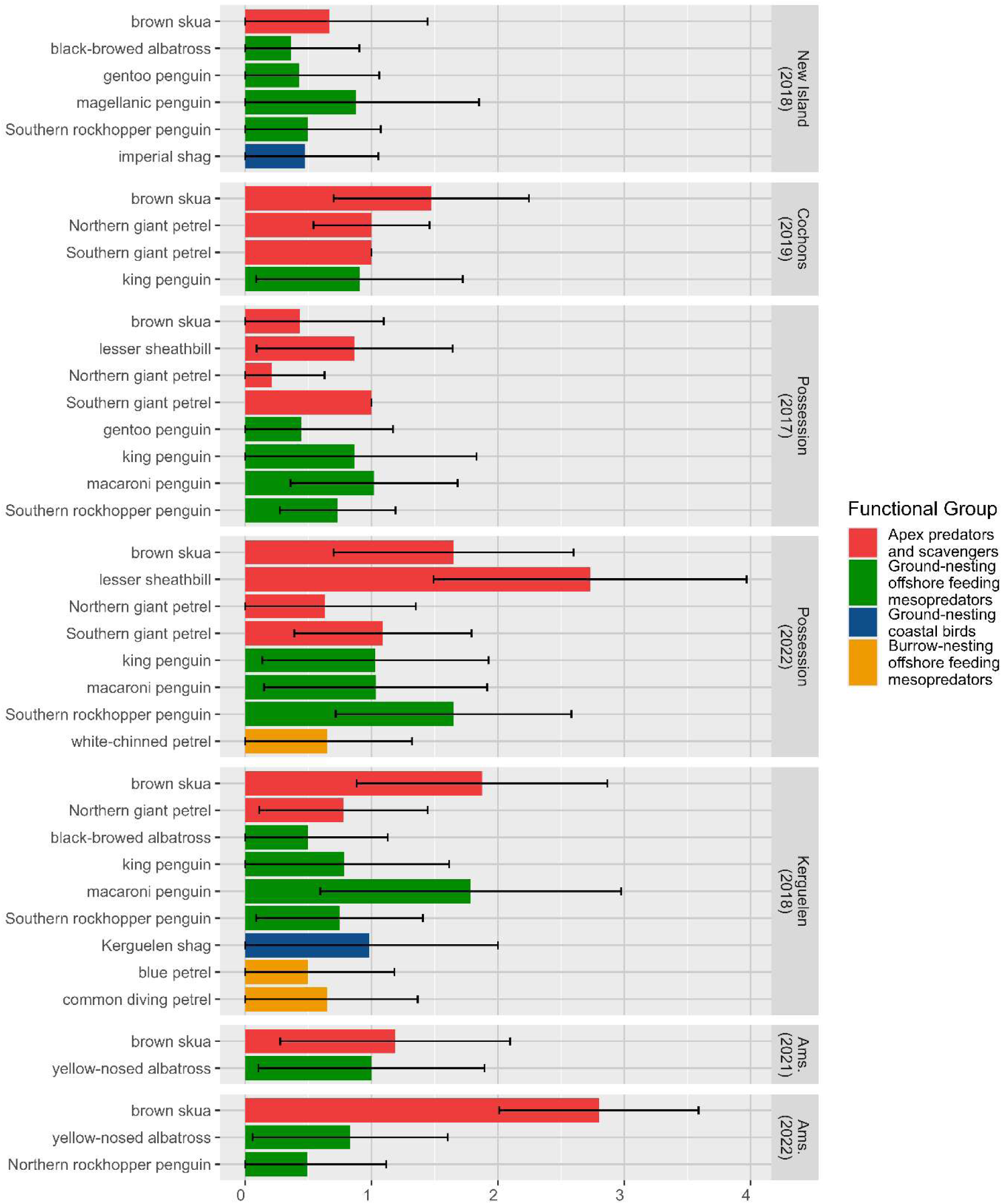
Diversity of infectious agents (here, specific richness) in seabird species from five different subantarctic areas. Brackets represent for standard deviation.

**Figure 4.**
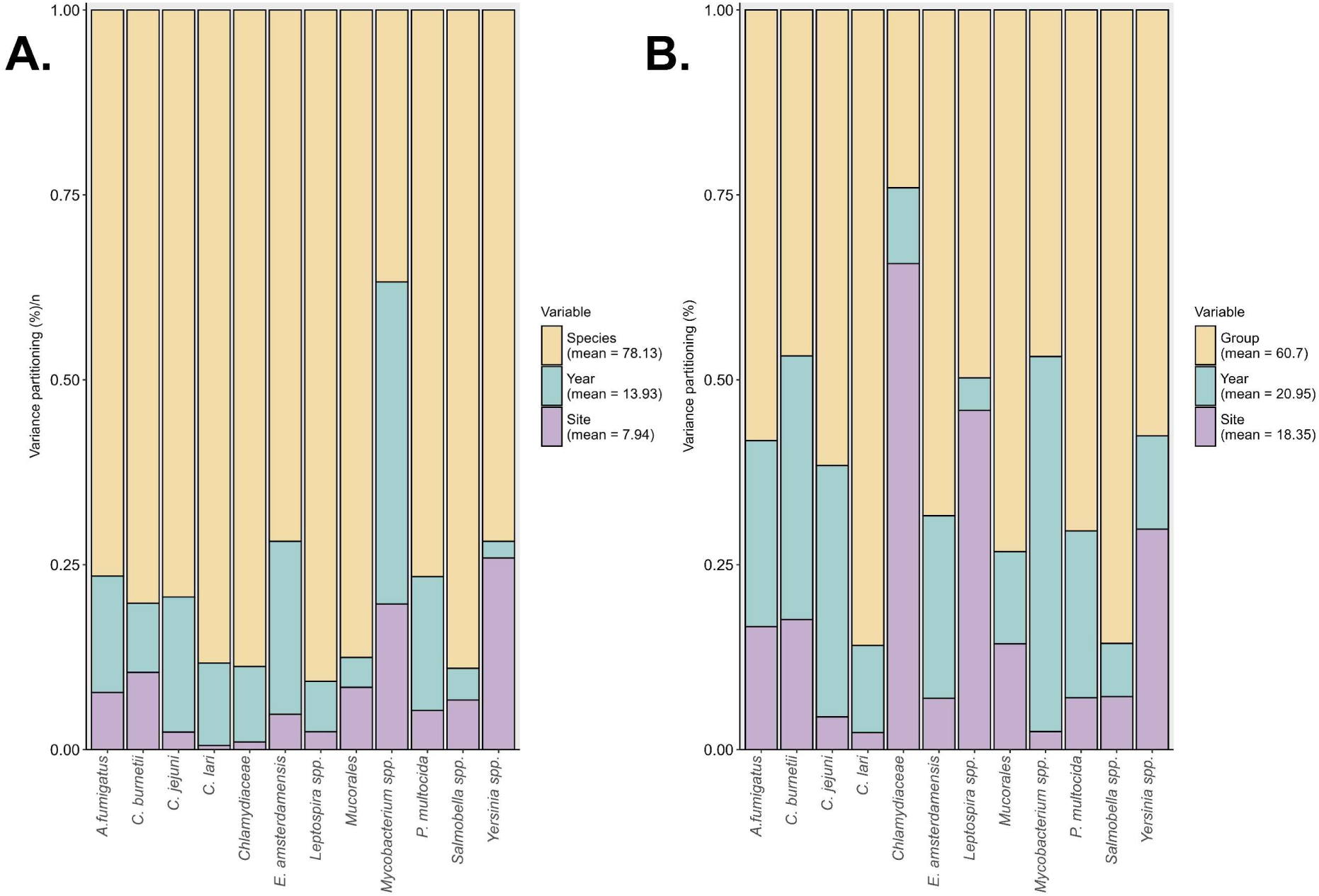
Variance partitioning among the fixed and random effects in island-scale models with species (A) and ecological group (B) as fixed effect. (respectively, Island_v6 and Island_v3 models). The height of each bar represents the proportion of variance explained by each variable for the different infectious agents.

### 2. JSDM convergence and model selection

Visual inspection of the β, Ɣ and Ω coefficients confirmed convergence for all JSDM models (**Table 1**). Running three series of models across three spatial scales enabled selecting three models based on WAIC, Tjur’s explanatory R² and AUC values: *Island_v6 (∼ host species + (1|WIS) + (1|year))*, *Regio_v6 (∼ host species + (1|island) + (1|year))* and *Global_v6 (∼ host species + (1|island) + (1|year)).* Satisfactory explanatory and predictive values indicated a good fit with datasets. The most prevalent IAs were those for which all models exhibited the best explanatory power and are hence the IAs for which we had the best predictive power. However, as investigating interactions among IA communities was an objective of this study, we retained IAs with low but non-zero prevalence in the dataset.

### 3. Meta-ecosystem and ecosystem scales: temporal and spatial variations

The effects of *WIS* and *year* on the presence of IA were assessed using an island-scale model on data from three POS sampling campaigns (**Table 1**). The best model (*Island_v6*) indicated that both factors had limited influence, with *WIS* explaining 7.9% and year explaining 13.9% of the model-explained variance (12.7%) in IA community (1,0 and 1,8% of total variance, respectively) (**Figure 4**). However, the results were heterogeneous: *WIS* explained up to 2,4% of variation for *Mycobacterium* spp. and 3,0% for *Yersinia* spp., whereas *year* accounted for the major part (6,4%) of the variation for *Mycobacterium* spp.

Comparing variance partitioning between *specie*s and *functional group* (**Figure 4**) showed that including species identity as a fixed effect markedly reduced the proportion of variance attributed to the *WIS* random effect. This suggests that the apparent site effect is largely driven by differences in host species differences rather than by local environmental conditions.

However, large-scale factors could not be assessed in the regional- and global-scale models because the sampling campaigns were not conducted in the same year, which prevented the detangling of *year* and *island* effects. Consequently, the predictive and explanatory values were similar across all models in each of the regional- and global-scale series (**Table 1**). We therefore retained both *year* and *island* as a single random effect and focused further analyses on local ecological factors.

### 4. Macro-community dimension: influence of host features

Using *host* s*pecies* as the explanatory variable produced greater explanatory power than using *functional group;* and the proportion of variation explained by the functional group was consistently lower. Including *host species* as a fixed effect improved the fit of the model, suggesting that features not captured by the *functional group* definition may play an important role.

Estimates of the β coefficients from the three selected models (*Island_v6, Regio_v6, Global_v6*) (**Figure 5**) showed that apex predators and scavengers were generally more infected by most of the studied IAs. Brown skuas exhibited positive β coefficients in all three models, and lesser sheathbills exhibited positive β values in the *Island_v6* model. By contrast, ground-nesting mesopredators such as penguins and albatrosses, especially burrowing white-chinned petrels (*Procellaria aequinoctialis*), exhibited negative β coefficients for seven out of 12 IAs in *Island_v6*, suggesting reduced IA infection in this species.

**Figure 5.**
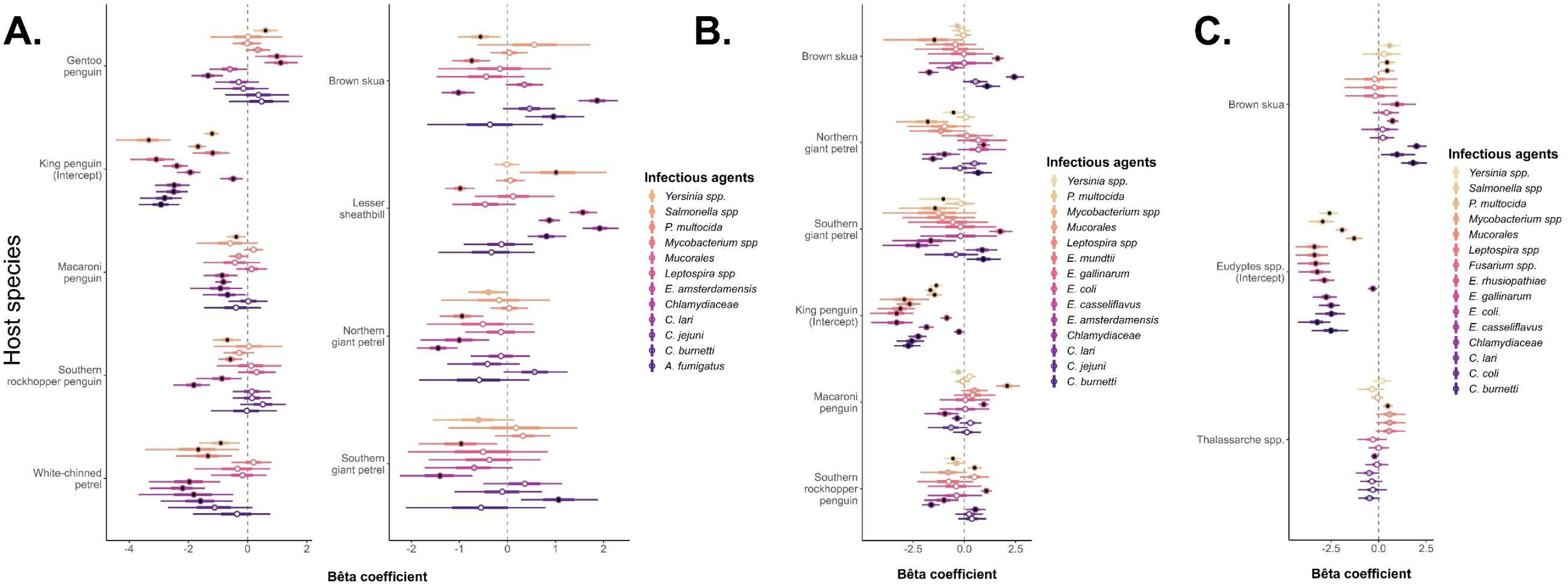
Estimation of beta coefficients for host species on the presence/absence each IA of a tested panel in the island-scale model (A), the regional-scale model (B), and the global-scale model (C). Each circle represents the mean and the bar the 95% credibility interval of the beta coefficient value distribution. A black circle indicates a credible difference from the intercept (PS ≥ 0.95), while a white circle indicates PS <0.95. Intercept values are derived from a comparison with the null model.

Among the IAs of particular interest (pathogenic or extensively studied in this system), *P. multocida* infection appeared largely homogeneous, with no significant differences between host species except for brown skuas in *Global_v6* (βmean = 0.44 [0.27-0.61], PS = 0.95). At both the island and regional scales, all species were less infected with *Chlamydiaceae* and *E. amsterdamensis* than the king penguins, while lesser sheathbills were more infected with these two IAs (0.87 [0.78-0.96], PS = 0.98 and 1.56 [1.46-1.69], PS = 1.0 in *Island_v6*). Conversely, at the regional scale (*Regio_v6*), all species were more infected with *E. coli* than king penguins.

A key feature of the JSDM is its ability to predict community characteristics, such as specific richness (PRS), from the β coefficients (**Supplementary Material S7**). PRS was not related to functional group (**Figure 3**), as gentoo penguins (*Pygoscelis papua*) and king penguins had the second and third highest PRS values (0.80 [0.35; 1.66] and 0.68 [0.26; 1.58], respectively), which exceeded those of brown skuas (0.64 [0.23; 1.52]) and giant petrels. The highest PRS value was observed in the lesser sheathbills (1.58 [0.76; 2.60]), while the lowest value was found in the burrowing white-chinned petrels (0.22 [0.02; 0.84]).

### 5. Infra-community dimension: influence of IAs features

Estimating the Ɣ coefficients provided insight into the relationship between host species and IA traits, particularly with regard to transmission mode. In the *Island_v6* model, burrowing white-chinned petrels (Ɣ mean = -1.44 [-1.75; -1.15], PS = 0.99) and, to a lesser extent, macaroni penguins (-0.44 [-0.61; -0.28], PS = 0.99) were less infected by directly transmitted IAs than other penguin species (**Figure 6**). No differences were detected for environmentally transmitted IAs in this model. However, in the *Regio_v6* model, macaroni penguins and the Northern giant petrels (*Macronectes halli*) were more (0.82 [0.53; 1.11], PS =0.95) or less (-1.18 [-1.64; - 0.76], PS =0.98) infected, respectively, than king penguins. Lesser sheathbills (0.61 [0.41; 0.79], PS = 0.95) and brown skuas (0.79 [0.56; 0.92], PS = 0.99) were more infected by directly-transmitted IAs in the *Island_v6* and *Global_v6* models, respectively.

**Figure 6.**
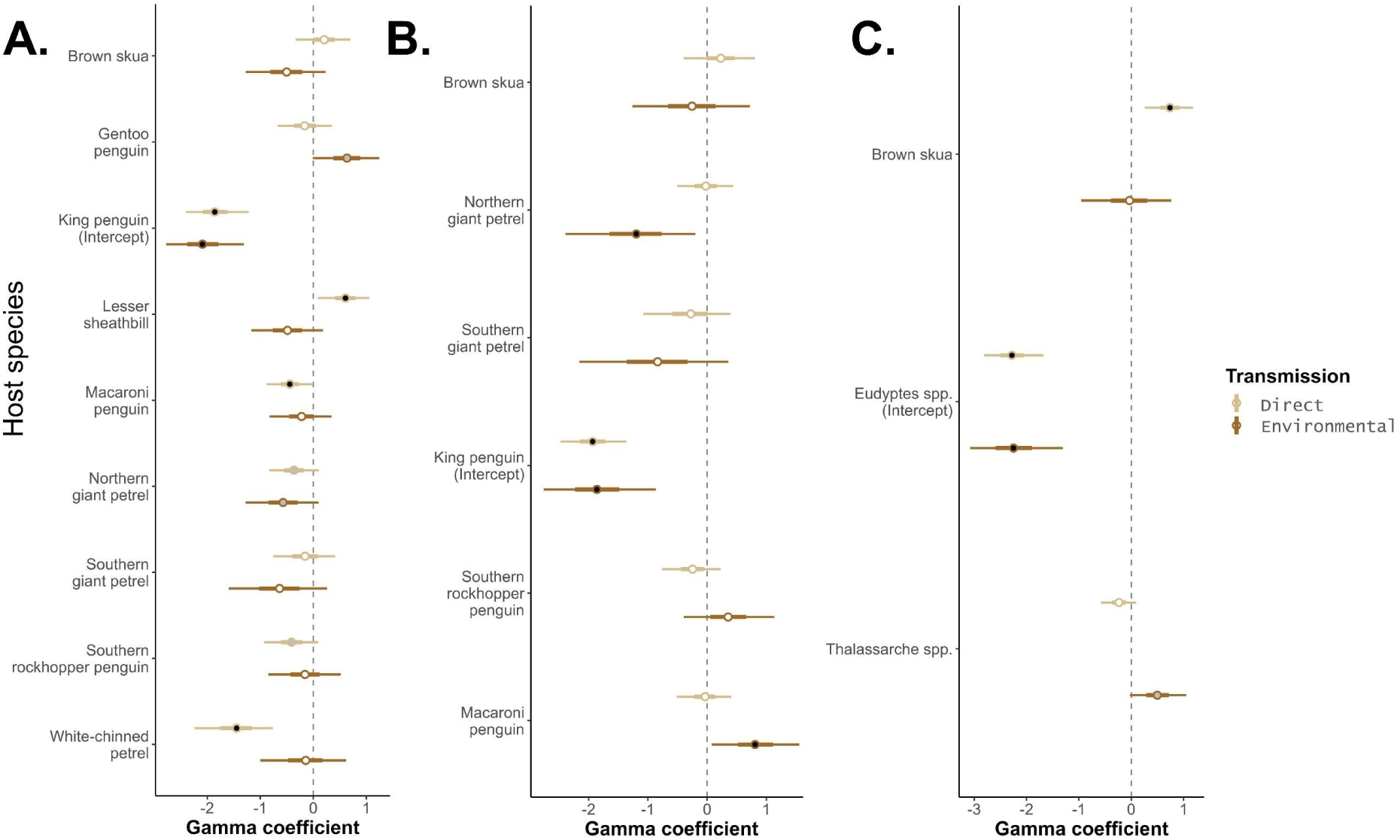
Estimation of gamma coefficients for the probability of host species becoming infected with directly- or environmentally-transmitted IAs in the island-scale model (A), regional-scale model (B), and the global-scale model (C). Each circle represents the mean and the bar the 95% credibility interval of the beta coefficient values distribution. A black circle indicates a credible difference from the intercept (PS ≥ 0.95), while a white circle indicates PS <0.95. Intercept values are derived from a comparison with the null model.

After controlling for intrinsic and extrinsic variables, as well as IA traits, the residual variance–covariance matrix (Ω) revealed potential associations between IAs. At the year scale, positive co-occurrence was observed among several directly transmitted IAs (**Figure 7.A**). Notably, *P. multocida* co-occurred positively with *Chlamydiaceae, E. amsterdamensis*, and *Mycobacterium* spp., suggesting that the prevalence of the former was highly correlated with the prevalence of the latter each year. At the *WIS* scale (**Figure 7.B**), positive co-occurrence was also observed between *Mycobacterium* sp. and *Yersinia* spp.

**Figure 7.**
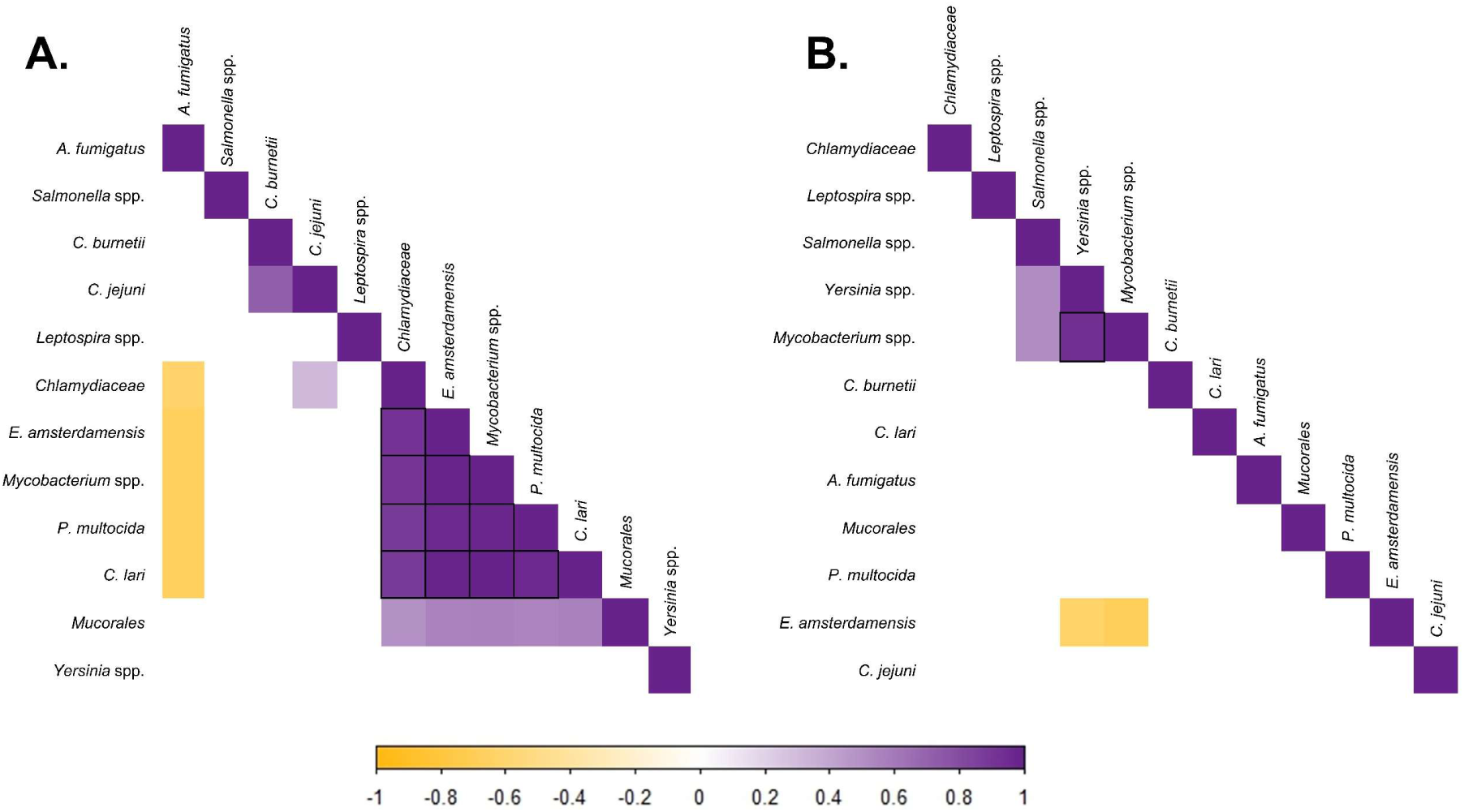
Associations between presences of infectious agents in seabirds after considering the effects of host species at the year level (A) or within-island site level (B) in the island-scale model. Purple and yellow indicate a positive or negative co-occurence, respectively (colors are only shown for values of posterior support greater than 0.75; boxed cells represent posterior values greater than 0.95).

## DISCUSSION

Understanding the patterns and drivers of infectious disease dynamics at different levels of biological organization is a key challenge in disease ecology. In this study, we examined IAs in seabird across five different subantarctic islands on a large scale. Using a hierarchical framework for modelling IA communities, we present new insights into how extrinsic and intrinsic drivers influence host-IA communities at meta-ecosystem, ecosystem, macro- community and infra-community dimensions.

### 1. Drivers at meta-ecosystem and ecosystem scales

As expected, IA communities were qualitatively similar across the subantarctic territories. Almost a fifth of the detected IAs were present on all islands, and around three quarters were present on at least two. This could reflect the similarity of bird communities and the climatic conditions that influence the large-scale distribution of IAs (Cohen *et al*., 2016).

Conducting a large-scale study also enabled us to document the widespread presence of IAs of interest for conservation or human health concerns, such as *P. multocida, Campylobacter* spp., and *Chlamydiaceae*. For instance, *C. lari* was found in almost all species, particularly brown skuas and penguins. These findings corroborate those of Cerdà-Cuéllar *et al*., (2019) concerning seabirds in the Southern Ocean. Similarly, the agent of avian cholera, *P. multocida*, was detected in all subantarctic territories. This aligns with a previous serological survey (Tornos *et al*., in prep), which reported seropositivity indicating exposure to *P. multocida* in seabirds from the same populations, even if seroprevalence was much lower in the Falkland Islands (11% of brown skuas) contrary to our PCR detection results. These results raise several eco-epidemiological questions: (1) Why do mortality events associated with avian cholera outbreaks only occur on certain islands, such as Amsterdam Island (Jaeger *et al*., 2018), Marion Island (Cooper *et al*., 2009) and the Antarctic Peninsula (Iervolino *et al*., 2025), when the pathogen appears to circulate across all subantarctic territories? (2) Could discrepancies between seroprevalence (Tornos *et al*., in prep) and our results using a direct detection method be explained by the circulation of different strains that escape detection of specific antibodies by existing serological assays? These findings highlight the need for further investigations including a more detailed characterization of the IAs (e.g. IA species, strain, pathotype and/or serotype). In particular, sequence analyses of specific IA could provide information to answer pressing issues in disease ecology such as dispersal processes and the evolution of large-scale strain lineages (Clessin *et al*., 2025).

At a lower spatial scale, the effect of the within-island site on the IA communities appears to be minimal, as no qualitative or quantitative differences were detected between the colonies. This is likely due to the high level of connectivity between colonies through seabirds movements while feeding, scavenging or prospecting (Boulinier, 2023; Lamb *et al*., 2023), which could facilitate IA dispersal and the homogenization of IA communities. However, it should be noted that our study only considered IAs that are directly or environmentally transmitted, and which can be transported through bird movements. Including avian vector-borne IA, notably tick-borne IA in this system (such as *Borrelia* spp. or flaviviruses) could have revealed a stronger *WIS* effect, as transmission would have been constrained by tick blood meals and limited tick mobility, potentially resulting in strong spatial structuring for such IA (Kada *et al*., 2017).

### 2. Drivers at macrocommunity dimension

Defining appropriate functional groups by integrating key host traits is essential for drawing robust conclusions in disease ecology. Here, we applied a principal component analysis to define the ‘candidate functional groups’ (Hooper *et al*., 2002), selecting the ecological traits most likely to influence IA exposure and circulation *a priori*. Specifically, we considered trophic level as a potential driver of interspecific IA circulation and breeding characteristics relevant for both intra- and interspecific circulation. However, at the macro-community dimension, these functional groups had less explanatory power than host species identity in the fitted models (**Figure 4**), which prevent us from identifying overarching drivers clearly. These results may be explained by several factors. Firstly, important factors such as host competence, susceptibility to IA and host phylogeny, as well as immune features, were probably overlooked among taxonomic groups (Fountain-Jones *et al*., 2018). Secondly, ecological traits are not always clear-cut. Albatrosses and white-chinned petrels, for example, are considered as mesopredators in our study but are increasingly being described as carrion scavengers at sea (Cherel & Klages, 1998; Schunck *et al*., 2025), which blurs their trophic classification. Thirdly, species may not always be the most relevant ecological unit through which to explore infectious eco-epidemiological processes, given that substantial within-species heterogeneity can occur in functional traits. Female giant petrels, for example, forage more at sea (’mesopredator traits’) whereas males are more competitive scavenging on land (González-Solís *et al*., 2000). This could result in sex-specific differences in IA exposure (Tornos *et al*., in prep). Finally, some traits are dynamic and species may shift functional groups over time (Jax, 2005). In our study system, both brown skuas and lesser sheathbills were assigned to the ‘terrestrial predators and scavengers’ group during the study period. However, skuas migrate over long distances during the winter (Delord, 2015), whereas lesser sheathbills are permanent residents, which could significantly influence IA transmission.

Moreover, simplifying a complex system by defining functional groups is further complicated in disease ecology, given that there is no uniform, non-specific response to a given function (Jax, 2005). Responses are likely to rely on a large variety of IA traits, such as host specificity, host barriers and transmission modes. In our study, we focused on transmission mode because we thought it would be the most important parameter to investigate in relation with the host functional groups of our system (Bralet *et al*., 2025). However, this approach could be complexified by considering the IA life cycle (Manlove *et al*., 2022) and host specificity. This would likely require a larger range of samples, to ensure sufficient sample size for the different values of each explanatory variable. Including additional transmission modes, particularly vector-borne IAs, could be of high interest (Berland *et al*., (2025). Looking for such IAs was however not possible with the type of sample we collected.

Indeed, host species was the strongest predictor of IA community composition among individual hosts, highlighting the importance of host species in shaping IA communities (Dallas *et al*., 2019).

### 3. Drivers at infra-community dimension

One objective of this study was to explore the relationship between host ecology and IA traits, particularly with regard to transmission mode. As burrows provide protected spaces and stable local microclimates (Retief *et al*., 2017; Kollath *et al*., 2020), they could potentially favor the long-term survival of IAs. Furthermore, burrowing behavior could increase exposure to soil-borne IAs. Therefore, we expected burrowing seabirds to be more exposed to environmentally transmitted IAs than other species. However, this was not observed, suggesting that these IAs are distributed relatively across the system. Interestingly, burrowing petrels were less frequently infected with directly transmitted IAs, likely because they directly fly between burrows and the sea (Warham, 1996), which limits contact - and thus transmission - with conspecifics (Ochoa-Sánchez *et al*., 2024) or other species at the colony site (but see Boulinier (2023) for discussion on at-sea transmission).

While IA associations should be interpreted as hypotheses rather than certainties due to the limitations of our dataset, our results highlight numerous positive co-occurrences within the IA community. However, these models did not allow us to determine the direction of causality. At the *WIS* scale, co-occurrence was observed between *Yersinia* spp. and *Mycobacterium* spp. As these IAs were characterized as mainly environmental, primarily belonging to the *M. terrae* complex in the latter case (see **Supplementary Material S8**), this primarily suggested the implication of environmental factors, such as humidity, temperature and vegetation, in shaping the distribution of these IAs. At the year scale, co-occurrences were primarily observed among directly transmitted IAs, including E. *amsterdamensis*, *P. multocida*, *C. lari*, and *Chlamydiaceae*. Such observations can be explained by year-driven exposure or year-driven susceptibility to IAs (Sweeny & Albery, 2022). The former corresponds, for example, to environmental conditions favoring such direct IAs or host population dynamics. The latter could be due to factors favouring individuals in poor body condition or to the sporadic circulation of an IA that triggers immunity at a population level in certain years, thereby favouring the circulation of such directly transmitted IAs. This highlights the need to combine such studies with environmental monitoring, as well as observation of mortality and morbidity, and exposure data on the serological status of populations.

## Conclusion

In this study, we present the results of a large-scale investigation of the prevalence of IA infections in seabirds across five different subantarctic islands. The majority of IAs were shared between several islands or present on all islands, notably the agent of avian cholera *P. multocida*, which raises concerns about risks for the health of subantarctic seabird populations. Within an island, no heterogeneity among sites was detected. Our findings call for further multi-dimension studies that would combine complementary approaches to disentangle the complex processes at play in the dynamics of hosts and IAs interactions.

## Statement of authorship

TBo, TBr, JT, AG, SM and KL conceived the study. TBo, AG, JT, ML, AMP, TBr and RA, AG performed field work or help for DNA extraction. Tbr and SM were responsible for IA specific PCR set development and validation and TBr, CG and RA performed the Htrt PCR. TBr, FB and JT carried out the statistical analyses and TBr produced the figures. TBr wrote the first version of the manuscript and all authors provided inputs.

## Acknowledgments

We thank Fabrice Le Bouard, Nicolas Giraud, Marine Bely, Aline Flechet, Corisande Abiven, Juliette Baron, Louisiane Burkart, Lucie Gauchet, Robin Dardel, Camille De Pasquale, Zorhia-Lys Guillerm for their help at various stages of sampling or laboratory work, and Lorraine Michelet, Mégane Gasque, Alexandre Dremeau, Guillaume Désoubeaux, Sabine Delannoy and the National Reference Centre for Antibiotic Resistance and the Associated Enterococcus Laboratory for their help in designing some of the PCR systems, providing positive controls and participating in the confirmation and analysis of results. Many thanks to Guy Baele for his help with proofreading and advice. We would like to thank Innovative Diagnostics for supplying us with extraction controls. This research was funded by the ANR ECOPATHS project (ANR-21-CE35-0016). We acknowledge support from the French Polar Institute (IPEV ECOPATH-1151), BiodivRestore REMOVE_DISEASE project (ANR-21-BIRE-0006), France relance TVACALBA project (NextGen Europe), Réserve Naturelle Nationale des Terres Australes Françaises, Zone Atelier Antarctique (ZATA), OSU OREME (ECOPOP), and CNRS INEE SEE-Life ECOPATH.

## Reference

Berland, F., Bourret, V., Peroz, C., Malandrin, L., Bonsergent, C., Bailly, X., Masseglia, S., Nouvel, L.-X., Lagrée, A.-C., Rouxel, C., Dimeglio, C., Izopet, J., Ollivier, V., Boulinier, T., Villena, I., Aubert, D., Sluydts, V., Loc’h, G.L., Bonnet, A., Chaval, Y., Merlet, J., Lourtet, B., Gilot-Fromont, E. & Verheyden, H. (2025) Effects of host sex, age and behaviour on co-infection patterns in a wild ungulate. Parasitology, 1–51.

Boulinier, T. (2023) Avian influenza spread and seabird movements between colonies. Trends in Ecology & Evolution, 38, 391–395.

Bralet, T., Aaziz, R., Tornos, J., Gamble, A., Clessin, A., Lejeune, M., Galon, C., Michelet, L., Lesage, C., Jeanniard du Dot, T., Desoubeaux, G., Guyard, M., Delannoy, S., Moutailler, S., Laroucau, K. & Boulinier, T. (2025) High-throughput microfluidic real-time PCR as a promising tool in disease ecology. Journal of Animal Ecology, 94, 1625–1637.

Bralet, T., Clessin, A., Lejeune, M., McMahon, C., Gamble, A., Tornos, J. & Boulinier, T. (in prep) Seabirds, pinnipeds and their infectious agents as model systems for addressing pressing questions in infectious disease ecology.

Caron, A., Cappelle, J., Cumming, G.S., de Garine-Wichatitsky, M. & Gaidet, N. (2015) Bridge hosts, a missing link for disease ecology in multi-host systems. Veterinary Research, 46, 83.

Caron, A., de Garine-Wichatitsky, M., Ndlovu, M. & Cumming, G.S. (2012) Linking avian communities and avian influenza ecology in southern Africa using epidemiological functional groups. Veterinary Research, 43, 73.

Cerdà-Cuéllar, M., Moré, E., Ayats, T., Aguilera, M., Muñoz-González, S., Antilles, N., Ryan, P.G. & González-Solís, J. (2019) Do humans spread zoonotic enteric bacteria in Antarctica? Science of The Total Environment, 654, 190–196.

Cherel, Y. & Klages, N. (1998) *A review of the food of albatrosses*. *Albatros biology and conservation*, Surrey Beatty and Sons, Chipping Norton, Australia.

Clessin, A., Briand, F.-X., Tornos, J., Lejeune, M., De Pasquale, C., Fischer, R., Souchaud, F., Hirchaud, E., Hong, S.L., Bralet, T., Guinet, C., McMahon, C.R., Grasland, B., Baele, G. & Boulinier, T. (2025) Circumpolar spread of avian influenza H5N1 to southern Indian Ocean islands. Nature Communications, 16, 8463.

Clucas, G.V., Younger, J.L., Kao, D., Emmerson, L., Southwell, C., Wienecke, B., Rogers, A.D., Bost, C.-A., Miller, G.D., Polito, M.J., Lelliott, P., Handley, J., Crofts, S., Phillips, R.A., Dunn, M.J., Miller, K.J. & Hart, T. (2018) Comparative population genomics reveals key barriers to dispersal in Southern Ocean penguins. Molecular Ecology, 27, 4680–4697.

Cohen, J.M., Civitello, D.J., Brace, A.J., Feichtinger, E.M., Ortega, C.N., Richardson, J.C., Sauer, E.L., Liu, X. & Rohr, J.R. (2016) Spatial scale modulates the strength of ecological processes driving disease distributions. Proceedings of the National Academy of Sciences of the United States of America, 113, E3359–3364.

Cooper, J., Crawford, R.J.M., De Villier, M.S., Dyer, B.M., Hofmeyr, G.J.G. & Jonker, A. (2009) Disease outbreaks among penguins at sub-Antarctic Marion Island: a conservation concern. Marine Ornithology, 193–196.

Dallas, T.A., Laine, A.-L. & Ovaskainen, O. (2019) Detecting parasite associations within multi-species host and parasite communities. Proceedings of the Royal Society B: Biological Sciences, 286, 20191109.

Daszak, P., Cunningham, A.A. & Hyatt, A.D. (2000) Emerging infectious diseases of wildlife--threats to biodiversity and human health. *Science (New York*, N.Y.), 287, 443–449.

Delord, K., Barbraud, C., Bost, C.-A., Cherel, Y., Guinet, C. & Weimerskirch, H. (2014) Atlas of top predators from French Southern Territories in the Southern Indian Ocean, CNRS.

Descamps, S., Forbes, M.R., Gilchrist, H.G., Love, O.P. & Bêty, J. (2011) Avian cholera, post-hatching survival and selection on hatch characteristics in a long-lived bird, the common eider *Somateria mollisima* . 39–48.

Devleesschauwer, B., Torgerson, P., Charlier, J., Levecke, B., Praet, N., Roelandt, S., Smit, S., Dorny, P., Berkvens, D. & Speybroeck, N. (2022) prevalence: Tools for Prevalence Assessment Studies.

Dias, M.P., Martin, R., Pearmain, E.J., Burfield, I.J., Small, C., Phillips, R.A., Yates, O., Lascelles, B., Borboroglu, P.G. & Croxall, J.P. (2019) Threats to seabirds: A global assessment. Biological Conservation, 237, 525–537.

Dobson, A.P., Pimm, S.L., Hannah, L., Kaufman, L., Ahumada, J.A., Ando, A.W., Bernstein, A., Busch, J., Daszak, P., Engelmann, J., Kinnaird, M.F., Li, B.V., Loch-Temzelides, T., Lovejoy, T., Nowak, K., Roehrdanz, P.R. & Vale, M.M. (2020) Ecology and economics for pandemic prevention. Science, 369, 379–381.

Elderd, B.D., Mideo, N. & Duffy, M.A. (2022) Looking across Scales in Disease Ecology and Evolution*. The American Naturalist.

Fecchio, A., Svensson-Coelho, M., Bell, J., Ellis, V.A., Medeiros, M.C., Trisos, C.H., Blake, J.G., Loiselle, B.A., Tobias, J.A., Fanti, R., Coffey, E.D., Faria, I.P.D., Pinho, J.B., Felix, G., Braga, E.M., Anciães, M., Tkach, V., Bates, J., Witt, C., Weckstein, J.D., Ricklefs, R.E. & Farias, I.P. (2017) Host associations and turnover of haemosporidian parasites in manakins (Aves: Pipridae). Parasitology, 144, 984–993.

Fernández-i-Marín, X. (2016) ggmcmc: Analysis of MCMC Samples and Bayesian Inference. Journal of Statistical Software, 70, 1–20.

Fountain-Jones, N.M., Hutson, K.S., Jones, M., Nowak, B.F., Turnbull, A., Younger, J., O’Reilly, M., Watkins, E., Guernier-Cambert, V., Cooley, L. & Hamede, R. (2024) One Health on islands: Tractable ecosystems to explore the nexus between human, animal, terrestrial, and marine health. *BioScience*, biae101.

Fountain-Jones, N.M., Pearse, W.D., Escobar, L.E., Alba-Casals, A., Carver, S., Davies, T.J., Kraberger, S., Papeş, M., Vandegrift, K., Worsley-Tonks, K. & Craft, M.E. (2018) Towards an eco-phylogenetic framework for infectious disease ecology. Biological Reviews, 93, 950–970.

Gamble, A., Bazire, R., Delord, K., Barbraud, C., Jaeger, A., Gantelet, H., Thibault, E., Lebarbenchon, C., Lagadec, E., Tortosa, P., Weimerskirch, H., Thiebot, J., Garnier, R., Tornos, J. & Boulinier, T. (2020) Predator and scavenger movements among and within endangered seabird colonies: Opportunities for pathogen spread. Journal of Applied Ecology, 57, 367–378.

González-Solís, J., Croxall, J.P. & Wood, A.G. (2000) Sexual dimorphism and sexual segregation in foraging strategies of northern giant petrels, Macronectes halli, during incubation. Oikos, 90, 390–398.

Gupta, P., Robin, V.V. & Dharmarajan, G. (2020) Towards a more healthy conservation paradigm: integrating disease and molecular ecology to aid biological conservation. Journal of Genetics, 99, 65.

Hooper, D.U., Solan, M., Symstad, A., Áz, S.D., Gessner, M.O., Buchmann, N., Degrange, V., Grime, P., Hulot, F., Mermillod-Blondin, F., Roy, J., Spehn, E. & Peer, L.V. (2002) Species diversity, functional diversity, and ecosystem functioning. Biodiversity and Ecosystem Functioning (ed. by M. Loreau), S. Naeem), and P. Lnchausti), pp. 195–208. Oxford University PressOxford.

Iervolino, M., Günther, A., Begeman, L., Aguado, B., Bestebroer, T.M., Bellido-Martin, B., Coerper, A., Fornillo, M.V., Fusaro, B., Ibañez, A.E., Leijten, L., Lisovski, S., Mañez, M.B., Reade, A., Run, P. van, Soto, F., Wallis, B., Dewar, M., Alcamí, A., Beer, M., Vanstreels, R.E.T. & Kuiken, T. (2025) The expanding avian influenza panzootic: skua die-off in Antarctica. 2025.04.25.650384.

Inchausti, P. & Weimerskirch, H. (2002) Dispersal and metapopulation dynamics of an oceanic seabird, the wandering albatross, and its consequences for its response to long-line fisheries. Journal of Animal Ecology, 71, 765–770.

Jaeger, A., Lebarbenchon, C., Bourret, V., Bastien, M., Lagadec, E., Thiebot, J.-B., Boulinier, T., Delord, K., Barbraud, C., Marteau, C., Dellagi, K., Tortosa, P. & Weimerskirch, H. (2018) Avian cholera outbreaks threaten seabird species on Amsterdam Island. PLOS ONE, 13, e0197291.

Jax, K. (2005) Function and “functioning” in ecology: what does it mean? Oikos, 111, 641–648.

Kada, S., McCoy, K.D. & Boulinier, T. (2017) Impact of life stage-dependent dispersal on the colonization dynamics of host patches by ticks and tick-borne infectious agents. Parasites & Vectors, 10, 375.

Knief, U., Bregnballe, T., Alfarwi, I., Ballmann, M.Z., Brenninkmeijer, A., Bzoma, S., Chabrolle, A., Dimmlich, J., Engel, E., Fijn, R., Fischer, K., Hälterlein, B., Haupt, M., Hennig, V., Herrmann, C., Veld, R. in ‘t, Kirchhoff, E., Kristersson, M., Kühn, S., Larsson, K., Larsson, R., Lawton, N., Leopold, M., Lilipaly, S., Lock, L., Marty, R., Matheve, H., Meissner, W., Morrison, P., Newton, S., Olofsson, P., Packmor, F., Pedersen, K.T., Redfern, C., Scarton, F., Schenk, F., Scher, O., Serra, L., Sibille, A., Smith, J., Smith, W., Sterup, J., Stienen, E., Strassner, V., Valle, R.G.p, Bemmelen, R.S.A. van, Veen, J., Vervaeke, M., Weston, E., Wojcieszek, M. & Courtens, W. (2024) Highly pathogenic avian influenza causes mass mortality in Sandwich Tern Thalasseus sandvicensis breeding colonies across north-western Europe. Bird Conservation International, 34, e6.

Kollath, D.R., Teixeira, M.M., Funke, A., Miller, K.J. & Barker, B.M. (2020) Investigating the Role of Animal Burrows on the Ecology and Distribution of Coccidioides spp. in Arizona Soils. Mycopathologia, 185, 145–159.

Kueffer, C., Drake, D.R. & Fernández-Palacios, J.M. (2014) Island biology: looking towards the future. Biology Letters, 10, 20140719.

Lamb, J., Tornos, J., Dedet, R., Gantelet, H., Keck, N., Baron, J., Bely, M., Clessin, A., Flechet, A., Gamble, A. & Boulinier, T. (2023) Hanging out at the club: Breeding status and territoriality affect individual space use, multi-species overlap and pathogen transmission risk at a seabird colony. Functional Ecology, 37, 576–590.

Lane, J.V., Jeglinski, J.W.E., Avery-Gomm, S., Ballstaedt, E., Banyard, A.C., Barychka, T., Brown, I.H., Brugger, B., Burt, T.V., Careen, N., Castenschiold, J.H.F., Christensen-Dalsgaard, S., Clifford, S., Collins, S.M., Cunningham, E., Danielsen, J., Daunt, F., D’entremont, K.J.N., Doiron, P., Duffy, S., English, M.D., Falchieri, M., Giacinti, J., Gjerset, B., Granstad, S., Grémillet, D., Guillemette, M., Hallgrímsson, G.T., Hamer, K.C., Hammer, S., Harrison, K., Hart, J.D., Hatsell, C., Humpidge, R., James, J., Jenkinson, A., Jessopp, M., Jones, M.E.B., Lair, S., Lewis, T., Malinowska, A.A., McCluskie, A., McPhail, G., Moe, B., Montevecchi, W.A., Morgan, G., Nichol, C., Nisbet, C., Olsen, B., Provencher, J., Provost, P., Purdie, A., Rail, J., Robertson, G., Seyer, Y., Sheddan, M., Soos, C., Stephens, N., Strøm, H., Svansson, V., Tierney, T.D., Tyler, G., Wade, T., Wanless, S., Ward, C.R.E., Wilhelm, S.I., Wischnewski, S., Wright, L.J., Zonfrillo, B., Matthiopoulos, J. & Votier, S.C. (2024) High pathogenicity avian influenza (H5N1) in Northern Gannets (*Morus bassanus*): Global spread, clinical signs and demographic consequences. Ibis, 166, 633–650.

Levin, S.A. (1992) The Problem of Pattern and Scale in Ecology: The Robert H. MacArthur Award Lecture. Ecology, 73, 1943–1967.

Locke, S.A., McLaughlin, J.D. & Marcogliese, D.J. (2013) Predicting the similarity of parasite communities in freshwater fishes using the phylogeny, ecology and proximity of hosts. Oikos, 122, 73–83.

Longmire, J., Maltbie, M. & Baker, R. (1997) Use of “lysis buffer” in DNA isolation and its implications for museum collections. Museum of Texas Tech University, 1–3.

Makowski, D., Ben-Shachar, M.S., Chen, S.H.A. & Lüdecke, D. (2019) Indices of Effect Existence and Significance in the Bayesian Framework. Frontiers in Psychology, 10.

Manlove, K., Wilber, M., White, L., Bastille-Rousseau, G., Yang, A., Gilbertson, M.L.J., Craft, M.E., Cross, P.C., Wittemyer, G. & Pepin, K.M. (2022) Defining an epidemiological landscape that connects movement ecology to pathogen transmission and pace-of-life. Ecology Letters, 25, 1760–1782.

Moon, K.L., Chown, S.L. & Fraser, C.I. (2017) Reconsidering connectivity in the sub-Antarctic. Biological Reviews, 92, 2164–2181.

Moss, W.E., McDevitt-Galles, T., Calhoun, D.M. & Johnson, P.T.J. (2020) Tracking the assembly of nested parasite communities: Using β-diversity to understand variation in parasite richness and composition over time and scale. Journal of Animal Ecology, 89, 1532–1542.

Ochoa-Sánchez, M., Acuña-Gómez, E.P., Moraga, C.A., Gaete, K., Acevedo, J., Eguiarte, L.E. & Souza, V. (2024) The ephemeral microbiota: Ecological context and environmental variability drive the body surface microbiota composition of Magellanic penguins across subantarctic breeding colonies. Molecular Ecology, 33, e17472.

Oksanen, J., Simpson, G.L., Blanchet, F.G., Kindt, R., Legendre, P., Minchin, P.R., O’Hara, R.B., Solymos, P., Stevens, M.H.H., Szoecs, E., Wagner, H., Barbour, M., Bedward, M., Bolker, B., Borcard, D., Borman, T., Carvalho, G., Chirico, M., Caceres, M.D., Durand, S., Evangelista, H.B.A., FitzJohn, R., Friendly, M., Furneaux, B., Hannigan, G., Hill, M.O., Lahti, L., Martino, C., McGlinn, D., Ouellette, M.-H., Cunha, E.R., Smith, T., Stier, A., Braak, C.J.F.T. & Weedon, J. (2025) vegan: Community Ecology Package.

Ovaskainen, O. & Abrego, N. (2020) Joint Species Distribution Modelling: With Applications in R, Cambridge University Press.

Ovaskainen, O., Tikhonov, G., Norberg, A., Guillaume Blanchet, F., Duan, L., Dunson, D., Roslin, T. & Abrego, N. (2017) How to make more out of community data? A conceptual framework and its implementation as models and software. Ecology Letters, 20, 561–576.

Pearce, J. & Ferrier, S. (2000) Evaluating the predictive performance of habitat models developed using logistic regression. Ecological Modelling, 133, 225–245.

Pertierra, L.R., Segovia, N.I., Noll, D., Martinez, P.A., Pliscoff, P., Barbosa, A., Aragón, P., Raya Rey, A., Pistorius, P., Trathan, P., Polanowski, A., Bonadonna, F., Le Bohec, C., Bi, K., Wang-Claypool, C.Y., González-Acuña, D., Dantas, G.P.M., Bowie, R.C.K., Poulin, E. & Vianna, J.A. (2020) Cryptic speciation in gentoo penguins is driven by geographic isolation and regional marine conditions: Unforeseen vulnerabilities to global change. Diversity and Distributions, 26, 958–975.

Plowright, R.K., Sokolow, S.H., Gorman, M.E., Daszak, P. & Foley, J.E. (2008) Causal inference in disease ecology: investigating ecological drivers of disease emergence. Frontiers in Ecology and the Environment, 6, 420–429.

Retief, L., Bennett, N.C., Jarvis, J.U.M. & Bastos, A.D.S. (2017) Subterranean Mammals: Reservoirs of Infection or Overlooked Sentinels of Anthropogenic Environmental Soiling? EcoHealth, 14, 662–674.

Sáez-Ventura, Á., López-Montoya, A.J., Luna, Á., Romero-Vidal, P., Palma, A., Tella, J.L., Carrete, M., Liébanas, G.M. & Pérez, J.M. (2022) Drivers of the Ectoparasite Community and Co-Infection Patterns in Rural and Urban Burrowing Owls. Biology, 11, 1141.

Schunck, F., Souza, R., Donadio, D.N., Correa, L., Bicudo, R., Souza, M.O.D., Silva, P.G.C.E., Dias, E., Legal, E. & Barata, F. (2025) White-chinned Petrel Procellaria aequinoctialis feeding on a dead dolphin. Marine Ornithology, 53, 201–203.

Selmi, S. & Boulinier, T. (2001) Ecological Biogeography of Southern Ocean Islands: The Importance of Considering Spatial Issues. The American Naturalist, 158, 426–437.

Stephens, P.R., Altizer, S., Smith, K.F., Alonso Aguirre, A., Brown, J.H., Budischak, S.A., Byers, J.E., Dallas, T.A., Jonathan Davies, T., Drake, J.M., Ezenwa, V.O., Farrell, M.J., Gittleman, J.L., Han, B.A., Huang, S., Hutchinson, R.A., Johnson, P., Nunn, C.L., Onstad, D., Park, A., Vazquez-Prokopec, G.M., Schmidt, J.P. & Poulin, R. (2016) The macroecology of infectious diseases: a new perspective on global-scale drivers of pathogen distributions and impacts. Ecology Letters, 19, 1159–1171.

Sweeny, A.R. & Albery, G.F. (2022) Exposure and susceptibility: The Twin Pillars of infection. Functional Ecology, 36, 1713–1726.

Talmon, I., Pekarsky, S., Bartan, Y., Thie, N., Getz, W.M., Kamath, P.L., Bowie, R.C.K. & Nathan, R. (2025) Using wild-animal tracking for detecting and managing disease outbreaks. Trends in Ecology & Evolution, 40, 760–771.

Tikhonov, G., Opedal, Ø.H., Abrego, N., Lehikoinen, A., de Jonge, M.M.J., Oksanen, J. & Ovaskainen, O. (2020) Joint species distribution modelling with the r-package Hmsc. Methods in Ecology and Evolution, 11, 442–447.

Tjur, T. (2009) Coefficients of Determination in Logistic Regression Models—A New Proposal: The Coefficient of Discrimination. The American Statistician, 63, 366–372.

Tompkins, D.M., Dunn, A.M., Smith, M.J. & Telfer, S. (2011) Wildlife diseases: from individuals to ecosystems: Ecology of wildlife diseases. Journal of Animal Ecology, 80, 19–38.

Tornos, J., Gamble, A., Clessin, A., Bralet, T., Lejeune, M., Connan, M., Ryan, P.G., Weimerskirch, H., Gonzalez-Solis, J., Catry, P., Barbraud, C., Delord, K., Viblanc, V.B., Flechet, A., Baron, J., Caza, F., Tounsi, A., Ferchiou, S., Perret, S., Bost, C.-A., Steinfurth, A., Le Net, R., Thiebot, J.-B., Lebouard, F., Lesage, C., St-Pierre, Y., Gantelet, H. & Boulinier, T. (in prep) Large-scale exposure of southern seabirds to avian cholera *Pasteurella multocida* : scavengers as sentinels at different spatial scales.

Warham, J. (1996) The Behaviour, Population Biology and Physiology of the Petrels, Academic Press.

Watanabe S. (2010) Comparison of DIC and WAIC in Neural Bayes Learning. IEICE Technical Report; IEICE Tech. Rep., 110, 79–84.

Worsley-Tonks, K.E.L., Escobar, L.E., Biek, R., Castaneda-Guzman, M., Craft, M.E., Streicker, D.G., White, L.A. & Fountain-Jones, N.M. (2020) Using host traits to predict reservoir host species of rabies virus. PLOS Neglected Tropical Diseases, 14, e0008940.

