## Supplementary Material for "Drivers of host-infectious agent community associations in seabirds from sub-Antarctic oceanic islands"

### Drivers of host-parasite community associations in seabirds from sub-Antarctic territories.
### - Supplementary Material -

**Supplementary Material S1 | Information on sampling sites.** The sites are named after their French names.

| | | |
|---|---|---|
| 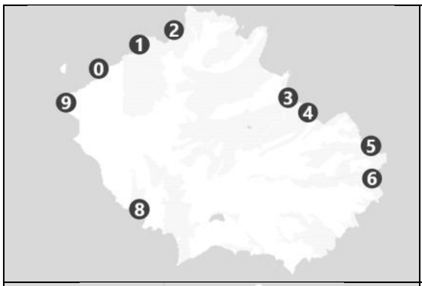 | **Possession Island,<br>Crozet Archipelago<br>(46°25'S, 51°45'E)** | 0 Mare aux éléphants<br>1 Champ des albatros<br>2 Jardin japonais<br>3 Baie américaine<br>4 Petite manchotière<br>5 Crique de la chaloupe<br>6 Baie du marin (scientific station)<br>7 Crique de Noël<br>8 La Pérouse<br>9 Les moines |
| 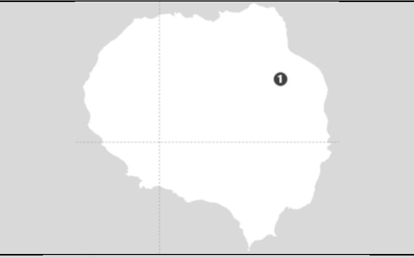 | **Cochons Island,<br>Crozet Archipelago<br>(49°28', 70°2')** | 1 Morne du Tamaris king penguin colony |
| 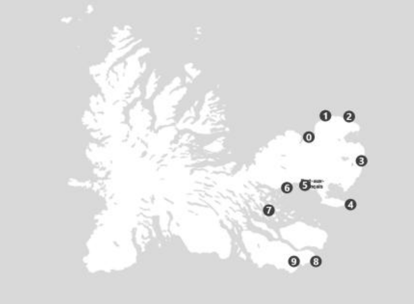 | **Kerguelen archipelago<br>(49°15'S, 69°10′E)** | 0 Cataracte<br>1 Cap Cotter<br>2 Cap noir<br>3 Ratmanoff<br>4 Pointe Suzanne<br>5 Port aux français (scientific station)<br>6 Cap Kidder<br>7 Mayes island<br>8 Sourcils noirs<br>9 Anse de l'Antarctique |
| 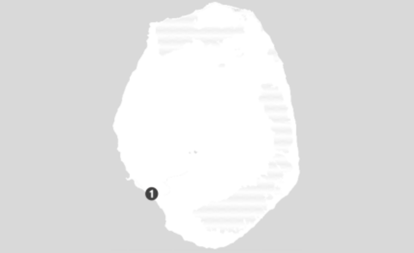 | **Amsterdam Island<br>(37°49′S, 77°33′E)** | 1 Entrecasteaux cliffs |
| 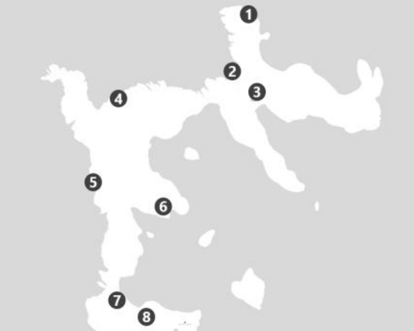 | **New Island,<br>Falkland Islands<br>(51°43′S, 61°18′W)** | 1 North bluff<br>2 Cathedral cliffs<br>3 Gentoo North<br>4 Landsend bluff<br>5 Rookery<br>6 Beef point<br>7 South harbour<br>8 Gentoo south |

**Supporting Informations – S2 | Prevalences and Clopper-Pearson 95% confidence intervals obtained simultaneously by Htrt PCR for each infectious agent depending on the species, sampling site and sample type (after validation step)**. Zero prevalences are not shown. Results are not shown in case of not found IA (*E. faecalis*, *E. hirae*, *Escherichia coli* STEC pathotype stx1, stx2 and stx2f, pathogenic *Leptospira* spp., *Mycobacterium tuberculosis* and *avium* complexes, pathogenic *Yersinia* spp, and *Toxoplasma gondii).*

| Island (year) | Species | Site | N | *A. fumigatus* | *Fusarium* spp. | *Mucorales* | *C. coli* | *C. jejuni* | *C. lari* | *Chlamydiaceae* | *Chlamydiifrater* spp. | *C. psittaci* | *C. burnetii* | *E. casseliflavus* | *E. faecium* | *E. gallinarum* | *E. mundtii* | *E. amsterdamensis* | *E. rhusiopathiae* | *E. coli* | *Leptospira* spp. | *Mycobacterium* spp. | *P. multocida* | *Salmonella* spp. | *Yersinia* spp. |
|---|---|---|---|---|---|---|---|---|---|---|---|---|---|---|---|---|---|---|---|---|---|---|---|---|---|
| New-Island (2018) | brown skua | Rookery South | 2 | | | | | | | | | | | | | | | | | 75,03 [35,92-100] | | | | | |
| | | Rookery C | 2 | | | | | | 52,12 [10,14-93,51] | | | | | | | | | | | 52,45 [9,91-94,25] | | 52,17 [10,03-93,91] | | | |
| | | South harbour | 3 | | | | | | | | | | | | | | | | | 42,17 [6,81-84,07] | | | | | |
| | | Beef point | 1 | | | | | | | | | | | | | | | | | | | | | | |
| | | Gentoo North | 10 | | | | | | | | | | | | | 17,45 [2,42-43,53] | | | | 52,63 [24,81-80,43] | | 35,08 [11,51-63,63] | | | |
| | | Gentoo South | 6 | | | | | | | | | | | | | | | | | 26,2 [3,66-61,51] | | 26,26 [3,97-60,83] | | | |
| | | Landsend bluff | 7 | | | | | | | | | | | | | | | | | 34,69 [9,16-67,79] | | | 35,06 [9-68,21] | | |
| | | Rookery bowl | 2 | | | | | | | | | | | | | | | | | 52,14 [9,8-94,11] | | | | | |
| | black-browed albatross | North bluff | 20 | | | | | | | | | | | | | | | | | 28,84 [12,1-50,17] | | | | | |
| | | Rookery A | 30 | | | | | | | | | | | | | | | | | 42,82 [26,06-61,1] | | | 9,8 [2,16-22,32] | | |
| | | Rookery C | 20 | | | | | | | | | | | | | 14,39 [3,3-32,11] | | | | 38,26 [18,82-59,73] | | 9,61 [1,2-24,85] | 14,28 [3,25-32,15] | | 9,58 [1,22-25,47] |
| | | Rookery E | 10 | | | | | | | | | | | | | | | | | | | | 17,53 [2,43-43,47] | | |
| | | Cathedral cliffs | 20 | | | | | | | | | | | | | | | | | 28,76 [11,67-50,09] | | | 9,59 [1,23-25,18] | | |
| | | Rookery B | 10 | | | | | | | | | | | | | | | | | 26,24 [6,44-53,79] | | | 17,68 [2,34-43,92] | | |
| | | Rookery face | 20 | | | | | | | | | | | | | | | | | 28,73 [11,86-49,63] | | | | | |
| | Southern rockhopper penguin | North bluff | 9 | | | | | | | | | | | | | 19,11 [2,7-47,11] | | | | 28,96 [7,18-58,79] | | | | | |
| | | Rookery South | 20 | | | | | | | | | | | | | | | | | 52,72 [31,32-73,83] | | | 14,28 [3,17-31,99] | | |
| | | Rookery A | 10 | | | | | | | | | | | | | | | | | 43,95 [17,75-73,11] | | | | | |
| | | Rookery C | 10 | | | | | | | | | | | | | | | | | 52,68 [24,55-80,35] | | | | | |
| | | Landsend bluff | 20 | | | | | | | 9,45 [1,19-24,75] | 9,4 [1,21-24,33] | | | | | | | | | 52,7 [31,3-73,96] | | 23,89 [8,67-43,99] | | | |
| | | Rookery bowl | 20 | | | | | | | | | | | | | | | | | 33,45 [15,22-55,22] | | | 14,38 [3,19-31,99] | | |
| | | Rookery E | 10 | | | | | | | | | | | | | | | | | 17,66 [2,37-44,08] | | | | | |
| | gentoo penguin | Gentoo North | 20 | | | 9,44 [1,25-24,55] | | | | 19 [5,87-38,26] | 9,53 [1,25-24,94] | | 9,48 [1,29-25,04] | | | | | | | 19,29 [5,84-38,58] | | | | | |
| | | Gentoo South | 20 | | 9,53 [1,18-24,9] | | | | | 14,16 [3,2-31,62] | 9,57 [1,29-24,98] | | | | | | | | | 23,85 [8,54-44,08] | | 9,59 [1,24-25,1] | 9,61 [1,2-25,3] | | |
| | imperial shag | North bluff | 10 | | | | | | | | | | | | | | | | | 26,09 [6,33-54,06] | | | 17,5 [2,4-43,1] | | |
| | | Rookery South | 20 | | | | | | 9,56 [1,26-24,8] | | | | | | | | | | | 9,66 [1,26-25,89] | | 9,64 [1,23-25,44] | 14,37 [3,17-32,07] | | |
| | | Rookery A | 20 | | | | | | | | | | | | | | | | | 38,3 [18,93-60,13] | | | 9,59 [1,3-25,03] | | |
| | | Rookery C | 20 | 9,6 [1,23-25,16] | | | | | 9,59 [1,21-25,61] | | | | | | | | | | | 57,43 [35,78-78,44] | 9,63 [1,23-25,23] | | 14,4 [3,13-32,09] | | |
| | Magellanic penguin | South harbour | 20 | | 14,36 [3,37-31,39] | | | | 9,56 [1,18-25,07] | 38,26 [18,95-59,97] | 33,44 [15,22-54,96] | | | | 9,61 [1,26-25,02] | | | | | 28,84 [12,01-50,07] | | 9,59 [1,27-25,27] | 23,88 [8,57-43,8] | | |
| | | Beef point | 5 | | | | | | | | | | | | | | | | | | | | 30,12 [4,6-67,15] | | |
| Cochons (2019) | brown skua | Morne du Tamaris | 19 | | | | | | 75,13 [53,41-92,96] | | | | | | | | | | | 70,17 [48,59-89,11] | | 10,01 [1,31-26,43] | | | |
| | king penguin | Morne du Tamaris | 63 | | | 3,22 [0,42-8,66] | | | 3,25 [0,39-8,83] | 42,11 [30,06-54,83] | 79,33 [67,92-89,5] | 8,1 [2,7-16,14] | | | | | | 8,06 [2,66-16] | | 24,34 [14,54-35,78] | | 8,07 [2,65-16] | 8,07 [2,78-15,99] | | 6,52 [1,83-14,08] |
| | Northern giant petrel | Morne du Tamaris | 20 | | | | | | 9,45 [1,26-24,39] | | | | | | | | | | | 89,85 [72,66-99,44] | | | 9,58 [1,2-24,95] | | |
| | Southern giant petrel | Morne du Tamaris | 8 | | | | | | | | | | | | | | | | | 89,93 [71,4-100] | | | | | |
| Possession (2017) | brown skua | Jardin japonais | 5 | | | | | | 30,2 [4,36-68,03] | | | | | | | | | | | | | | | | |
| | | Petite manchotière | 2 | | | | | | | | | | | | | | | | | | | | | | |
| | | Baie du marin | 3 | | | | | | 42,34 [7,2-84,75] | | | | | | | | | | | | | | | | |
| | | Baie américaine | 4 | | | | | | 35 [5,73-74,61] | | | | | | | | | | | 52,43 [15,34-89,43] | | | 35,27 [5,44-76,15] | | |
| | | La Pérouse | 6 | | | | | | | | | | | | | | | | | 39,39 [10,36-73,9] | | | | | |
| | | Champ des albatros | 3 | | | | | | 42,07 [7,47-84,74] | | | | | | | | | | | 41,9 [6,73-84,9] | | | | | |
| | king penguin | Jardin japonais | 30 | | | | | | | 36,22 [20,49-54,08] | 36,19 [20,32-53,82] | | | | | | | 6,53 [0,84-17,42] | | 19,72 [7,77-35,53] | | 32,77 [17,25-50,6] | 9,91 [2,22-22,64] | | 42,8 [25,64-61,25] |
| | | Baie américaine | 30 | 6,57 [0,85-17,91] | | | | | | 19,67 [7,82-35,54] | 9,9 [2,18-22,45] | 9,89 [2,16-22,53] | | | | | | | | 16,49 [5,72-31,27] | | 6,61 [0,83-17,67] | 9,79 [2,1-22,15] | | |
| | macaroni penguin | Jardin japonais | 29 | | | | | | | 13,59 [3,83-27,87] | 16,99 [6,07-32,38] | | | | | | | | | 61,08 [42,67-78,39] | 6,79 [0,85-18,15] | 6,7 [0,85-18,03] | | | 20,31 [8,05-36,84] |
| | | val austere | 20 | | | | | | | | | | | | | | | | | 85,8 [66,68-98,43] | 9,62 [1,24-25,01] | | 19,11 [5,68-38,24] | | 14,3 [3,25-31,97] |
| | Southern rockhopper penguin | Jardin japonais | 15 | | | | | | | | | | | | | | | | | 74,35 [50,31-93,69] | | | | | |
| | Northern giant petrel | Jardin japonais | 2 | | | | | | | | | | | | | | | | | 52,49 [9,91-94,48] | | | | | |
| | | Petite manchotière | 6 | | | | | | | | | | | 26,42 [4,06-61,46] | | | | | | | | | | | |
| | | Baie du marin | 4 | | | | | | | | 35,34 [5,23-76,67] | | | | | | | | | | | | | | |
| | | Baie américaine | 2 | | | | | | | | | | | | | | | | | | | | | | |
| | | Champ des albatros | 5 | | | | | | | | | | | | | | | | | 44,9 [12,26-81,44] | | | | | |
| | lesser sheathbill | Jardin japonais | 10 | | | | | | 43,89 [17,77-72,8] | | | | | | | | | | | | | | | | |
| | | Baie du marin | 10 | | | | | | 44,01 [17,76-72,96] | 44,04 [17,81-72,82] | 44,02 [17,5-73,13] | | | | | | | 26,31 [6,26-54,67] | | 26,21 [6,45-54,22] | | | | | |
| | | Baie américaine | 10 | | | | | | | 44 [17,81-73,12] | 34,94 [11,29-63,65] | | | | | | | 52,63 [24,76-80,88] | | | | | | | 17,83 [2,46-44,33] |
| | Southern giant petrel | Baie du marin | 1 | | | | | | | | | | | | | | | | | 66,57 [21,99-100] | | | | | |
| | | Baie américaine | 1 | | | | | | | | | | | | | | | | | 67,13 [22,79-100] | | | | | |
| | gentoo penguin | La Pérouse | 9 | | | | | | | | | | | | | | | | | 19,26 [2,59-47,13] | | | | | 38,39 [12,95-69,04] |

| | | | | | | | | | | | | | | | | | | | | | | | | |
|---|---|---|---|---|---|---|---|---|---|---|---|---|---|---|---|---|---|---|---|---|---|---|---|---|
| Possession (2022) | brown skua | Baie du marin | 10 | | | | | 44 [17,7-73,34] | | | | | | | | | 17,79 [2,36-44,26] | | 78,56 [50,54-97,73] | | 17,58 [2,47-43,78] | 17,42 [2,43-43,33] | | |
| | | Baie américaine | 10 | | | | | 61,36 [32,35-87,18] | 17,49 [2,4-43,48] | 17,58 [2,34-43,59] | | 43,93 [17,93-72,82] | | | | | | | 85,95 [61,88-99,3] | | | | | 26,45 [6,34-54,31] |
| | | La Pérouse | 10 | | | | | 61,26 [31,83-87,63] | | 17,4 [2,5-42,98] | | 17,79 [2,46-43,94] | | | | | | | 78,35 [49,96-97,65] | | | | | |
| | | Champ des albatros | 10 | | | | 35,01 [11,47-64,62] | 43,87 [17,59-72,74] | | 17,62 [2,45-43,52] | | | | | | | | | 44,11 [17,69-72,87] | | 26,45 [6,19-54,97] | 17,66 [2,43-43,71] | | |
| | king penguin | Jardin japonais | 20 | | | | 9,59 [1,19-25,11] | | 81,33 [61-96,41] | 57,29 [36-77,85] | 9,56 [1,24-25,13] | | | | | | | | | 9,56 [1,29-25,08] | 43,05 [23,45-64,24] | | | 38,25 [18,86-60,13] |
| | | Mare aux éléphants | 20 | | | | 9,68 [1,23-25,56] | | 67,04 [45,6-86,33] | 71,92 [50,9-89,59] | | | | | | | 28,68 [11,91-49,44] | | 9,47 [1,24-24,79] | | | | | 19,18 [5,88-38,31] |
| | | Baie de la chaloupe | 20 | | | | | 9,42 [1,27-24,26] | 47,86 [27,21-69,74] | 23,94 [8,69-43,73] | 14,23 [3,15-31,4] | | | | | | | | 19,25 [5,73-38,5] | | 9,6 [1,2-24,91] | | | |
| | | Petite manchotière | 20 | | | | | | 33,38 [15,32-54,75] | 19,13 [5,8-38,21] | 9,56 [1,21-25,06] | 9,54 [1,24-24,75] | | | | | | | 33,48 [15,4-54,9] | | 28,71 [11,89-49,36] | 9,55 [1,24-25,36] | | 9,56 [1,28-24,57] |
| | | Baie du marin | 17 | | | | | | 33,04 [13,87-55,93] | 38,86 [18,35-62,42] | | | | | | | | | 22,21 [7,03-43,17] | | 11,09 [1,46-28,98] | 11,08 [1,42-28,65] | | |
| | macaroni penguin | Jardin japonais | 20 | | | | | | 19,26 [5,82-38,44] | 23,77 [8,78-43,99] | | | | | | | | | 38,3 [19,1-59,54] | 9,56 [1,24-25,27] | 14,4 [3,26-31,87] | 19 [5,71-37,82] | | 14,33 [3,26-32,03] |
| | | Moines | 20 | | | | | | 14,29 [3,15-31,98] | 14,26 [3,22-31,83] | | | | | | | | | 47,77 [27,09-69,2] | | 14,5 [3,16-32,55] | 14,3 [3,18-31,66] | | |
| | | La Pérouse | 20 | | | | | | 14,26 [3,26-31,88] | 14,22 [3,22-31,62] | | 9,57 [1,26-25,33] | | | | 9,6 [1,28-25,12] | | | 85,81 [66,76-98,42] | | 23,81 [8,63-43,79] | 19,19 [5,73-38,37] | | 9,55 [1,2-25,07] |
| | Southern rockhopper penguin | Jardin japonais | 20 | | | | 14,38 [3,12-31,94] | 9,56 [1,16-25,13] | | | | 9,65 [1,27-24,87] | | | | | | | 90,01 [73,34-99,38] | 14,25 [3,23-31,51] | 38,34 [19,14-60,25] | | | 14,24 [3,24-31,32] |
| | Northern giant petrel | Baie du marin | 10 | | | | | | 17,71 [2,35-44,56] | | | | 17,66 [2,39-43,2] | | 17,58 [2,41-43,6] | | | | 35,11 [11,44-64,51] | | | 17,58 [2,46-43,91] | | 17,65 [2,4-43,51] |
| | | Baie américaine | 10 | | | | | | 17,59 [2,41-43,99] | 26,25 [6,22-54,17] | | 26,27 [6,38-54,4] | | | | | | | 52,78 [24,88-80,62] | | | 17,68 [2,28-43,63] | | 17,57 [2,42-43,68] |
| | | Champ des albatros | 10 | | | | | | | 17,55 [2,48-43,13] | | | | | | | | | | | | 17,52 [2,43-43,81] | | |
| | lesser sheathbill | Jardin japonais | 35 | | | | 14,23 [4,89-27,51] | 17,03 [6,68-31,2] | 90,7 [77,75-99,09] | 82,46 [67,51-94,6] | 8,57 [1,88-19,82] | | 5,66 [0,71-15,22] | | | | 53,93 [37,5-70,35] | | 59,78 [42,84-75,99] | | 8,48 [1,84-19,93] | 11,37 [3,34-23,53] | 8,52 [1,79-19,55] | 14,26 [4,81-27,73] |
| | | Baie du marin | 33 | | | | 15,03 [5,18-28,61] | 27,2 [13,59-43,73] | 87,23 [72,85-97,86] | 81,22 [65,43-93,9] | 18,02 [7,12-32,32] | | | | 6 [0,75-16,04] | | 42,23 [25,89-59,64] | | 75,05 [58,61-89,25] | 6,01 [0,77-16,18] | | 9,07 [1,9-20,91] | | 9,03 [1,96-20,69] |
| | | Baie américaine | 32 | | | | 21,55 [9,51-37,24] | 46,39 [29,74-63,9] | 95,87 [86,23-99,88] | 77,41 [60,84-91,22] | 15,44 [5,47-29,68] | | | | 6,24 [0,77-16,77] | | 55,79 [38,66-72,86] | | 68,09 [50,47-83,96] | | 12,35 [3,5-25,5] | 12,3 [3,62-25,18] | | 15,49 [5,31-29,8] |
| | white-chinned petrel | Barrage | 20 | | | | | | | | | | | | | | | | 28,58 [11,94-48,76] | | 38,23 [19,37-59,16] | | | 9,61 [1,22-24,88] |
| | Southern giant petrel | Baie du marin | 3 | | | | | 42,19 [7,15-84,8] | | | | 41,96 [7,25-85] | | | | | | | 79,7 [46,42-100] | | | 41,83 [6,74-84,7] | | |
| | | Baie américaine | 3 | | | | | | | | | | | | | | | | 62,48 [19,95-96,14] | | | | | |
| | | Champ des albatros | 5 | | | | | | | | | 29,84 [4,45-66,97] | | | | | | | 59,76 [23,53-92,07] | | | | | |
| Kerguelen (2018) | brown skua | Mayes | 4 | | | | | 35,11 [5,41-75,82] | | | | | | | | | | | 69,15 [28,34-97,35] | | | | | |
| | | Ratmanoff | 7 | | | | | 58,8 [26,21-88,61] | | | | | | | | | | | 88,97 [68,94-100] | | 23,47 [3,39-55,61] | | | 23,37 [3,32-55,28] |
| | | Cap Cotter | 11 | | | | | 80,59 [54,31-98,11] | 16,26 [2,24-40,77] | | | 16,35 [2,29-40,76] | | | | | | | 87,11 [63,16-99,43] | | 24,16 [5,86-50,29] | | | 16,39 [2,23-41,78] |
| | | Cataractes | 2 | | | | | 52,07 [10,32-93,72] | | | | | | | | | | | 75,29 [37,06-100] | | | 52,02 [9,95-93,93] | | |
| | blue petrel | Mayes | 30 | | | | | | | | | 6,55 [0,82-17,42] | | | | | | | 29,59 [15,01-46,91] | | 9,83 [2,14-22,46] | | | 6,54 [0,84-17,34] |
| | king penguin | Ratmanoff | 30 | | | | | | 36,18 [20,34-53,72] | 26,17 [12,37-43,38] | | | | | | | | | 26,24 [12,52-43,33] | | 6,5 [0,81-17,33] | 16,47 [5,64-31,75] | | 13,14 [3,78-26,97] |
| | | anse antarctique | 25 | | 7,73 [1-20,18] | | | | 23,46 [9,52-41,69] | 19,6 [6,83-36,93] | | | | | | | | | 35,23 [18,12-54,79] | | | 7,81 [0,99-20,76] | | 15,48 [4,68-31,3] |
| | macaroni penguin | Cap Cotter | 50 | | 90,77 [79,69-98,72] | | | 10,08 [3,39-20,04] | 54,54 [40,45-68,31] | 40,52 [27,17-54,57] | | | | | | | | | 52,67 [38,6-66,77] | 4,04 [0,51-10,88] | | 8,04 [2,28-16,76] | | 8,15 [2,33-17,26] |
| | | Sourcils noirs | 20 | | | | | | 23,85 [8,77-44,31] | 19,22 [5,69-38,54] | | | | | | | | | 62,16 [40,53-82,24] | | 9,44 [1,26-24,65] | | | 9,65 [1,27-25,35] |
| | black-browed albatross | Sourcils noirs | 30 | | 6,59 [0,83-17,5] | | | 6,58 [0,83-17,78] | | | | | | | | | | | 32,93 [17,35-50,82] | 6,59 [0,83-17,63] | 9,83 [2,2-22,2] | | | 6,45 [0,84-16,87] |
| | Southern rockhopper penguin | Mayes | 30 | | | | | | 6,51 [0,82-17,43] | 6,55 [0,82-17,52] | | | | | | | | | 52,69 [35,2-70,3] | | 13,13 [3,9-26,84] | | | 6,57 [0,82-17,48] |
| | | Cap kidder | 15 | | | | | 24,79 [7,49-47,81] | 12,46 [1,69-31,17] | 12,51 [1,64-32,28] | | | | | | | | | 36,98 [16,05-61,56] | | 18,34 [4,29-39,39] | 12,33 [1,65-31,89] | | |
| | | Cataractes | 30 | | | | | | 6,63 [0,88-17,68] | 9,9 [2,12-22,61] | | | | | | | | | 69,02 [51,31-84,93] | | 9,83 [2,18-22,11] | 6,56 [0,86-17,42] | | |
| | Kerguelen shag | Cataractes | 15 | | | | | 31,15 [11,69-55,08] | | | | | | | | | | | 18,61 [4,23-40,23] | | 12,46 [1,66-31,78] | | | 18,68 [4,24-40,85] |
| | | Cap noir | 15 | | | | | 12,43 [1,59-31,88] | | | | | | | | | | | 18,77 [4,41-40,79] | | 18,7 [4,16-41,44] | 18,51 [4,18-39,87] | 18,55 [4,24-40,16] | 43,24 [20,46-67,59] |
| | | pointe Suzanne | 30 | | | | | 49,27 [31,89-67,46] | | | | 13,22 [3,8-27,03] | | | | | | | 23,06 [10,14-39,15] | | 26,33 [12,73-42,9] | 9,94 [2,12-22,93] | | 13,19 [3,89-27,15] |
| | Northern giant petrel | Ratmanoff | 4 | | | | | 35,07 [5,46-75,01] | | | | | | | | | | | 69,33 [29,7-97,35] | | | | | |
| | | Cap Cotter | 1 | | | | | | | | | | | | | | | | 66,46 [22,04-100] | | | | | |
| | | port aux francais | 4 | | | | | | | | | | | | | | | | 52,58 [15,28-89,52] | | | | | |
| | common diving petrel | Mayes | 23 | | | | | 8,5 [1,11-22,73] | | | | 8,39 [1,06-22,17] | | | | | | | 37,8 [19,82-58,08] | | 21,09 [7,55-39,59] | | | |
| Ams. (2021) | yellow-nosed albatross | Entrecasteaux | 6 | 26,13 [3,82-60,22] | | | | | | | | | | | | | | | 39,23 [10,51-74,61] | | 39,51 [10,56-74,43] | | | 26,23 [3,7-60,68] |
| | brown skua | plateau des tourbieres | 5 | | | | | | | | | | | | | | | | 59,82 [22,96-92,16] | | | 45,02 [12,38-81,76] | | 29,93 [4,36-67,25] |
| | | Mare aux éléphants | 1 | | | | | 67,08 [22,78-100] | | | | | | | | | | | 66,75 [22,63-100] | | | | | |
| | | Entrecasteaux (plage) | 10 | | | | | 43,91 [17,7-73,03] | | | | 17,57 [2,36-43,63] | | | | | | | 61,34 [32,34-87,56] | | | | | |
| Ams. (2022) | yellow-nosed albatross | Entrecasteaux | 89 | 2,33 [0,29-6,27] | | | | | | | | 2,31 [0,28-6,37] | 2,3 [0,29-6,23] | | | | | | 33,47 [23,96-43,82] | | 48,64 [38,06-59,55] | 2,3 [0,29-6,38] | | 2,31 [0,29-6,37] |
| | brown skua | Entrecasteaux (plage) | 10 | | | 17,64 [2,35-43,91] | | 61,49 [32,84-87,64] | | | | 78,72 [51,52-97,89] | | | | | | 17,56 [2,32-43,68] | 91,64 [76,07-100] | | | 26,35 [6,35-54,43] | | |
| | Northern rockhopper penguin | Entrecasteaux | 75 | | | | | | | | | 5,44 [1,54-11,69] | 2,76 [0,35-7,68] | | | | | | 31,5 [21,39-42,75] | | 12,23 [5,82-20,42] | 4,1 [0,9-9,62] | 2,76 [0,34-7,5] | |

**Supplementary Material S3 | PCR sets added to the PCR assay of Bralet et al., 2025**. All probes were labelled with the FAM fluorophore and BHQ1 quencher.

| Infectious agent | Target | Name | Sequences (5'-3') | Lenght (bp) | Reference** | Reference genomes (GenBank accession number) | Positive contrôle |
|---|---|---|---|---|---|---|---|
| ***Chlamydiifrater*** | 16S | ChlFrater_16S_F<br>ChlFrater_16S_R<br>ChlFrater_16S_P | CTGGTACGCTCAAGTTTTCTG<br>GCGTGTGTGATGAAGGCTTTA<br>ATATTAGCCAAAATCCCTTATTCCCAGGCGAA | 90 | Adapted from Vorimore et al., (2021) | LR777654, OZ022385, LR777658 | culture (strains from Vorimore et al., (2021) |
| ***Enterococcus casseliflavus*** | pEM | EnteroCass_pEM_F<br>EnteroCass_pEM_R<br>EnteroCass_pEM_P | CCAATTCTTTATGAGGCAGACC<br>GGCGAAGTGATCAATCCAGC<br>AGCGCGGGATTATCCTGATGAGTTTATTC | 145 | This study | CP119296, CP116026, CP141640, CP046123, LR607377 | culture* |
| ***Enterococcus faescium*** | pEM | EnteroFaes_pEM_F<br>EnteroFaes_pEM_R<br>EnteroFaes_pEM_P | GTCCAATTGAGGTGTTCTTACC<br>CTC CGA TTC CTA GAA CAT TAG C<br>CAGCGTGTTTCGATCGGGATACGTC | 122 | This study | CP145124, CP131522, CP131520, CP144273, CP137476 | culture* |
| ***Enterococcus fecalis*** | pEM | EnteroFeca_pEM_F<br>EnteroFeca_pEM_R<br>EnteroFeca_pEM_P | TTGAGATCGATGTGTATTTGTGTG<br>CAGCTATTTCCTGATATTGATGAC<br>CTGAATAGGCACTTCTTTAATTATCGGCGAAG | 165 | This study | CP039296, CP019512, CP124898, CP110072 | culture* |
| ***Enterococcus gallinarum*** | sodA | EnteroGall_sodA_F<br>EnteroGall_sodA_R<br>EnteroGall_sodA_P | CTGAAGACATCAAAACAGCTGTC<br>GCTGCCAAATGTTTCTTCGATG<br>CGTAATAACGGTGGTGGTCATGCAAATCAC | 140 | This study | AJ387940, CP091204, AM490318, CP050816, EU021334, CP050485, CP116511, CP046307, AJ387915 | culture* |
| ***Enterococcus hirae*** | sodA | EnteroHir_sodA_F<br>EnteroHir_sodA_R<br>EnteroHir_sodA_P | TATTGATAAGCTAATGCAAGCGC<br>CTTTGCTTTACCCAATGTTTGTC<br>AAACCAGCAAATATGGGTTCAAGTGTAGGGAT | 112 | Adapted from Prichula et al., (2016) | CP055232, CP065992, CP072891, CP109807, AP027299, CP055229, CP055230, CP055231 | culture* |
| ***Enterococcus mundtii*** | sodA | EnteroMund_sodA_F<br>EnteroMund_sodA_R<br>EnteroMund_sodA_P | CAGACATGGATGCTATTCCATC<br>GCCATGATTTTCCAGAAGAATGA<br>TGGTTCGCATGTCCACCACCATTATTACG | 94 | Adapted from Prichula et al., (2016) | AP019810, AJ387918, JX436496, AP013036, CP083695, OV996158, CP022340, CP018061, CP025473, CP029066 | culture* |
| ***Escherichia coli*** | cdgR | cdgR-F<br>cdgR-R<br>cdgR-Taq | GCTATTTCCTGCCGATAAGAGA<br>CCAGGCAAAGAGTTTATGTTGA<br>ACCAGCAAGACCTTGTCGGCGTTGAG | / | Delannoy et al., (2020) | / | culture* |
| ***Escherichia coli STEC*** | stx1 | vt1-F<br>vt1-R<br>vt1-taq | GGATAATTTGTTTGCAGTTGATGTC<br>CAAATCCTGTCACATATAAATTATTTCGT<br>CCGTAGATTATTAAACCGCCCTTCCTCTGGA | / | Nielsen and Andersen 2003 | / | culture* |
| | stx2 | stx-F<br>stx-R<br>stx2-Taq | TTTGTYACTGTSACAGCWGAAGCYTTACG<br>CCCCAGTTCARWGTRAGRTCMACRTC<br>TCGTCAGGCACTGTCTGAAACTGCTCC | / | Perelle et al., 2004 | / | culture* |
| | stx2f | vtx2f-F<br>vtx2f-R<br>vtx2f-Taq | TGGGAAGCGAATACAATC<br>ATGGAATTAGCAGAAAAGAGACCG<br>TCTTACTGAACCAAACCAATAACAGGG | / | Delannoy et al., unpublished | / | culture* |
| ***16S*** | 16S | 16Snew-F1<br>16Snew-R1<br>16Snew-Taq | TGGAGCATGTGGTTTAATTCGA<br>TGCGGGACTTAACCCAACA<br>CACGAGCTGACGACARCCATGCA | / | Delannoy et al, (2023) | / | / |

*Culture of *Enterococcus* spp. and *E. coli* were provided by the National Reference Centre for Antibiotic Resistance and the Associated Enterococcus Laboratory (Laboratoire Associé Entérocoques, Caen, 14033, France) and COLiPATH Unit & Genomics Platform IdentyPath (Laboratory for Food Safety, ANSES, 94700 Maisons-Alfort, France), respectively.

**References:


- Delannoy S, Schouler C, Souillard R, Yousfi L, Le Devendec L, Lucas C, Bougeard S, Keita A, Fach P, Galliot P, Balaine L, Puterflam J, Kempf I. Diversity of Escherichia coli strains isolated from day-old broiler chicks, their environment and colibacillosis lesions in 80 flocks in France. Vet Microbiol. 2021 Jan;252:108923. doi: 10.1016/j.vetmic.2020.108923. Epub 2020 Nov 7. PMID: 33221068
- Lindsey RL, Garcia-Toledo L, Fasulo D, Gladney LM, Strockbine N. Multiplex polymerase chain reaction for identification of Escherichia coli, Escherichia albertii and Escherichia fergusonii. J Microbiol Methods. 2017 Sep;140:1-4. doi: 10.1016/j.mimet.2017.06.005. Epub 2017 Jun 6. PMID: 28599915; PMCID: PMC5603207.

- Nielsen EM, Andersen MT. Detection and characterization of verocytotoxin-producing Escherichia coli by automated 5' nuclease PCR assay. J Clin Microbiol. 2003 Jul;41(7):2884-93. doi: 10.1128/JCM.41.7.2884-2893.2003. PMID: 12843017; PMCID: PMC165313.
- Perelle, S., Dilasser, F., Grout, J., and Fach, P. (2004). Detection by 5′-nuclease PCR of Shiga-toxin producing *Escherichia coli* O26, O55, O91, O103, O111, O113, O145 and O157:H7, associated with the world's most frequent clinical cases. *Mol. Cell. Probes* 18, 185–192. doi: 10.1016/j.mcp.2003.12.004
- Delannoy S, Hoffer C, Tran ML, Madec JY, Brisabois A, Fach P, Haenni M. High throughput qPCR analyses suggest that Enterobacterales of French sheep and cow cheese rarely carry genes conferring resistances to critically important antibiotics for human medicine. Int J Food Microbiol. 2023 Oct 16;403:110303. doi: 10.1016/j.ijfoodmicro.2023.110303. Epub 2023 Jun 24. PMID: 37384974.
- Prichula, J., Pereira, R. I., Wachholz, G. R., Cardoso, L. A., Tolfo, N. C. C., Santestevan, N. A., Medeiros, A. W., Tavares, M., Frazzon, J., d'Azevedo, P. A., & Frazzon, A. P. G. (2016). Resistance to antimicrobial agents among enterococci isolated from fecal samples of wild marine species in the southern coast of Brazil. Marine Pollution Bulletin, 105(1), 51–57. https://doi.org/10.1016/j.marpolbul.2016.02.071
- Vorimore, F., Hölzer, M., Liebler-Tenorio, E. M., Barf, L.-M., Delannoy, S., Vittecoq, M., Wedlarski, R., Lécu, A., Scharf, S., Blanchard, Y., Fach, P., Hsia, R. C., Bavoil, P. M., Rosselló-Móra, R., Laroucau, K., & Sachse, K. (2021). Evidence for the existence of a new genus Chlamydiifrater gen. nov. inside the family Chlamydiaceae with two new species isolated from flamingo (Phoenicopterus roseus): Chlamydiifrater phoenicopteri sp. nov. and Chlamydiifrater volucris sp. nov. Systematic and Applied Microbiology, 44(4), 126200. https://doi.org/10.1016/j.syapm.2021.126200

**Supplementary Material S4 | List of infectious agents targeted by Htrt PCR in this study**

*Brucella* spp., *Campylobacter coli*, *C. jejuni*, *C. lari*, *Chlamydiaceae*, *Chlamydia psittaci*, *Chlamydiifrater* spp., *Coxiella burnetii*, *Enterococcus casseliflavus*, *E. faecalis*, *E. faecium*, *E. gallinarum*, *E. hirae*, *E. mundtii*, *Erysipelothrix amsterdamensis*, *E. rhusiopathiae*, *Escherichia coli*, *E. coli* STEC stx1, stx2 and stx2f pathotypes, *Leptospira spp*, pathogenic *Leptospira* spp., *Mycobacterium* spp., *M. tuberculosis* and *avium* complexes, *Pasteurella multocida*, *Salmonella* spp., *Yersinia* spp. and pathogenic *Yersinia* spp, *Aspergillus fumigatus*, *Fusarium* spp., Mucorales and *Toxoplasma gondii*.

**Supplementary Material S5 | Host species grouping according to ecological features using Principal Component Analysis (PCA).**

PCA was performed using the same life history trait data and script as described by Tornos et al. (2025). Information on the life history traits of seabird species from the Falkland Islands (which were not studied in Tornos et al., 2025) was added (extracted from Wilman et al., 2014; Cherel et al., 2005; Verheyden et al., 1991; Clausen et al., 2002; Thomson et al., 1993; Küpfer et al., 2023).

The results of a PCA, including several traits of the 18 species, showed that the species are spread along two main gradients explaining respectively 44,8% and 21,5% of the variance (**Table S4.1**, **Figure S4.1**). The first axis separates two groups of species: the terrestrial apex predators and scavengers, partially feeding on birds and mammals and marine or shore predators, feeding mostly on marine resources (fishes and/or crustaceans). The second axis separates three groups within the marine and shore predators: the ground nesting species that feed mainly on shore, the ground nesting species feeding at sea and the burrow nesting species feeding at sea.

According to these results, we distinguished four ecological groups of species (**Figure S4.2**): (i) the terrestrial apex predators and scavengers (skuas, giant petrels and sheathbills), (ii) the ground nesting coastal birds (gulls and shags), (iii) the ground nesting mesopredators (penguins and albatrosses and (iv) the burrow nesting mesopredators (small petrels). Each group is represented by a color for this study, respectively (i) red, (ii) blue, (iii) green and (iv) yellow.

**Table S4.1:** Proportion of variance explained by each axis and cumulative variance explained for the Principal Component Analysis of species traits.

| | Eigen value | Variance (%) | Cumulative variance (%) |
|---|---|---|---|
| **Dim.1** | 4.4832351 | 44.8 | 44.8 |
| **Dim.2** | 2.1524658 | 21.5 | 66.4 |
| **Dim.3** | 1.4884959 | 14.9 | 81.2 |
| **Dim.4** | 0.9766520 | 9.8 | 91.0 |
| **Dim.5** | 0.4369965 | 4.7 | 95.4 |

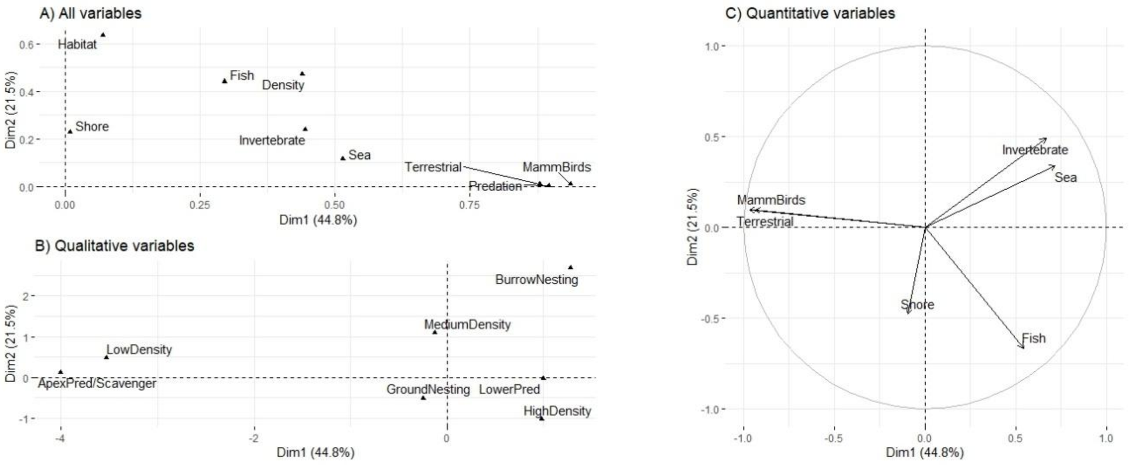


**Figure S2.1**: Coordinates in Dim.1 and Dim.2 of variables on correlation graph. A) all variables (absolute coordinates); B) qualitative variables and C) quantitative variables.

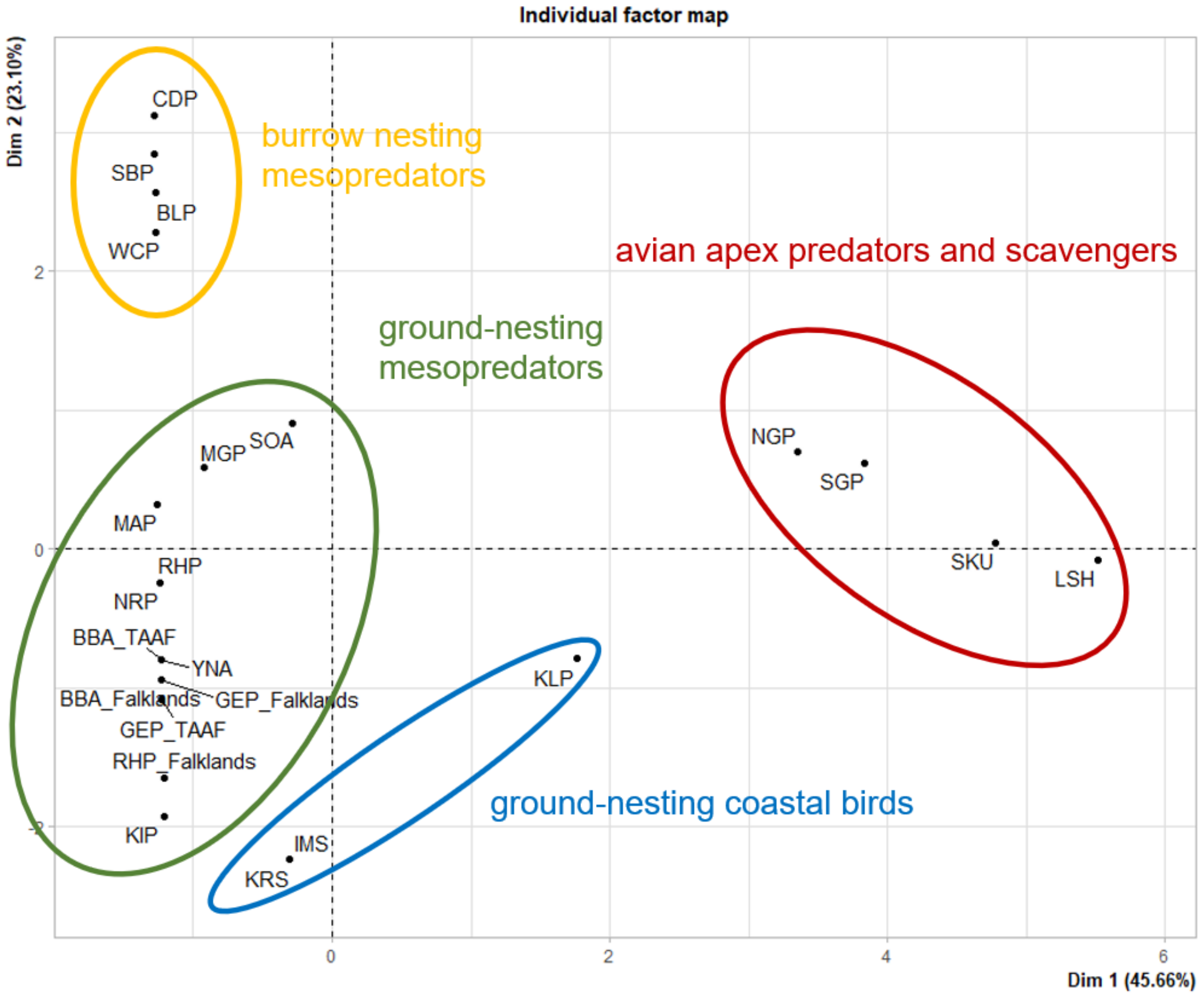


**Figure S2.2**: All species coordinates in correlation graph from the PCA analysis. Colors are relatives to the groups defined by the 2 main axes. *BBA: black-browed albatross, BLP: blue petrel, CDP: common diving petrel, GEP: gentoo penguin, IMS: imperial shag, KIP: king penguin, KLP: kelp gull, KRS: Kerguelen shag, LSH: lesser sheathbill, MAP: macaroni penguin, MGP: Magellanic penguin, NGP: Northern giant petrel, NRP: Northern rockhopper penguin, RHP: Southern rockhopper penguin, SBP: slender-billed prion, SGP: Southern giant petrel, SKU: brown skua, SOA: sooty albatross, WCP: white-chinned petrel, YNA: yellow-nosed albatross. TAAF: French Southern and Antarctic Lands*

#### Supplementary Material S6 | Classification of IA according to its main transmission mode (direct versus environmentally transmitted).

| Infectious agent | Main transmission mode | References* |
|---|---|---|
| ***A. fumigatus*** | environment | Arné et al., (2021) |
| ***Fusarium* spp.** | environment | Rheeder et al., (1990) |
| **Mucorales** | environment | Howard, (2002) |
| ***T. gondii*** | environment | Zaki et al., (2024) |
| ***Brucella* spp.** | direct | Wareth et al., (2020) |
| ***C. coli*** | direct | Cerdà-Cuéllar et al., (2019) |
| ***C. jejuni*** | direct | Cerdà-Cuéllar et al., (2019) |
| ***C. lari*** | direct | Cerdà-Cuéllar et al., (2019) |
| ***Chlamydiaceae*** | direct | Aaziz et al., (2015) |
| ***C. burnetti*** | direct | Loureiro et al., (2024) |
| ***E. casseliflavus*** | direct | Prichula et al., (2016) |
| ***E. faecalis*** | direct | Prichula et al., (2016) |
| ***E. faecium*** | direct | Prichula et al., (2016) |
| ***E. gallinarum*** | direct | Prichula et al., (2016) |
| ***E. hirae*** | direct | Prichula et al., (2016) |
| ***E. mundtii*** | direct | Prichula et al., (2016) |
| ***E. amsterdamensis*** | direct | Wang et al., (2010) |
| ***E. rhusiopathiae*** | direct | Wang et al., (2010) |
| ***E. coli*** | direct | Cardoso et al., (2023) |
| ***Leptospira* spp** | environment | Goarant et al., (2019; Samrot et al., (2021) |
| ***Mycobacterium* spp.** | environment | Bralet et al. (2025) |
| ***P. multocida*** | direct | Jaeger et al., (2020) |
| ***Salmonella* spp.** | direct | Tizard, (2004) |
| ***Yersinia* spp.** | direct | Falcão, (2014), Bralet et al. 2025 |

*References:


- Aaziz, R., Gourlay, P., Vorimore, F., Sachse, K., Siarkou, V. I., & Laroucau, K. (2015). Chlamydiaceae in North Atlantic Seabirds Admitted to a Wildlife Rescue Center in Western France. Applied and Environmental Microbiology, 81(14), 4581–4590. https://doi.org/10.1128/AEM.00778-15
- Arné, P., Risco-Castillo, V., Jouvion, G., Le Barzic, C., & Guillot, J. (2021). Aspergillosis in Wild Birds. Journal of Fungi, 7(3), 241. https://doi.org/10.3390/jof7030241
- Cardoso, M. D., Gonçalves, V. D., Grael, A. S., Pedroso, V. M., Pires, J. R., Travassos, C. E. P. F., Domit, C., Vieira-Da-Motta, O., Dos Prazeres Rodrigues, D., & Siciliano, S. (2023). Detection of Escherichia coli and other Enterobacteriales members in seabirds sampled along the Brazilian coast. Preventive Veterinary Medicine, 218, 105978. https://doi.org/10.1016/j.prevetmed.2023.105978
- Cerdà-Cuéllar, M., Moré, E., Ayats, T., Aguilera, M., Muñoz-González, S., Antilles, N., Ryan, P. G., & González-Solís, J. (2019). Do humans spread zoonotic enteric bacteria in Antarctica? Science of The Total Environment, 654, 190–196. https://doi.org/10.1016/j.scitotenv.2018.10.272
- Falcão, J. P. (2014). YERSINIA | Introduction. In C. A. Batt & M. L. Tortorello (Eds.), Encyclopedia of Food Microbiology (Second Edition) (pp. 831–837). Academic Press. https://doi.org/10.1016/B978-0-12-384730-0.00362-1
- Goarant, C., Trueba, G., Bierque, E., Thibeaux, R., Davis, B., & De La Pena-Moctezuma, A. (2019). Leptospira and Leptospirosis. In Michigan State University, J. B. Rose, B. Jiménez Cisneros, & UNESCO - International Hydrological Programme (Eds.), Water and Sanitation for the 21st Century: Health and Microbiological Aspects of Excreta and Wastewater Management (Global Water Pathogen Project). Michigan State University. https://doi.org/10.14321/waterpathogens.26

- Howard, D. H. (Ed.). (2002). Pathogenic Fungi in Humans and Animals (2nd ed.). CRC Press. https://doi.org/10.1201/9780203909102
- Jaeger, A., Gamble, A., Lagadec, E., Lebarbenchon, C., Bourret, V., Tornos, J., Barbraud, C., Lemberger, K., Delord, K., Weimerskirch, H., Thiebot, J.-B., Boulinier, T., & Tortosa, P. (2020). Impact of Annual Bacterial Epizootics on Albatross Population on a Remote Island. EcoHealth, 17(2), 194–202. https://doi.org/10.1007/s10393-020-01487-8
- Loureiro, F., Cardoso, L., Matos, A., Matos, M., & Coelho, A. C. (2024). Coxiella burnetii in wild birds from Europe. 14(4).
- Prichula, J., Pereira, R. I., Wachholz, G. R., Cardoso, L. A., Tolfo, N. C. C., Santestevan, N. A., Medeiros, A. W., Tavares, M., Frazzon, J., d'Azevedo, P. A., & Frazzon, A. P. G. (2016). Resistance to antimicrobial agents among enterococci isolated from fecal samples of wild marine species in the southern coast of Brazil. Marine Pollution Bulletin, 105(1), 51–57. https://doi.org/10.1016/j.marpolbul.2016.02.071
- Rheeder, J. P., Marasas, W. F. O., & Van Wyk, P. S. (1990). Fusarium species from Marion and Prince Edward Islands: Sub-Antarctic. South African Journal of Botany, 56(4), 482–486. https://doi.org/10.1016/S0254-6299(16)31045-6
- Samrot, A. V., Sean, T. C., Bhavya, K. S., Sahithya, C. S., Chan-drasekaran, S., Palanisamy, R., Robinson, E. R., Subbiah, S. K., & Mok, P. L. (2021). Leptospiral Infection, Pathogenesis and Its Diagnosis—A Review. Pathogens, 10(2), Article 2. https://doi.org/10.3390/pathogens10020145
- Tizard, I. (2004). Salmonellosis in wild birds. Seminars in Avian and Exotic Pet Medicine, 13(2), 50–66. https://doi.org/10.1053/j.saep.2004.01.008.
- Wang, Q., Chang, B. J., & Riley, T. V. (2010). Erysipelothrix rhusiopathiae. Veterinary Microbiology, 140(3–4), 405–417. https://doi.org/10.1016/j.vetmic.2009.08.012
- Wareth, G., Kheimar, A., Neubauer, H., & Melzer, F. (2020). Susceptibility of Avian Species to Brucella Infection: A Hypothesis-Driven Study. Pathogens, 9(2), 77. https://doi.org/10.3390/pathogens9020077
- Zaki, L., Olfatifar, M., Ghaffarifar, F., Eslahi, A. V., KarimiPourSaryazdi, A., Taghipour, A., Hamidianfar, N., Badri, M., & Jokelainen, P. (2024). Global prevalence of Toxoplasma gondii in birds: A systematic review and meta-analysis. Parasite Ep

**Supplementary Material – S7 | Relationship between the host species and the specific richness of infectious agents by the HMSC model at Island-scale**. Year and within-island sites were fixed as the more probable values in the dataset. Dots represent observed data. Each circle represents the predicted specific richness and the bar the 95% confidence interval of the estimate. Colors represents ecological groups : Red for terrestrial apex predators and scavengers, green for ground-nesting mesopredators and yellow for  burrow-nesting mesopredators.

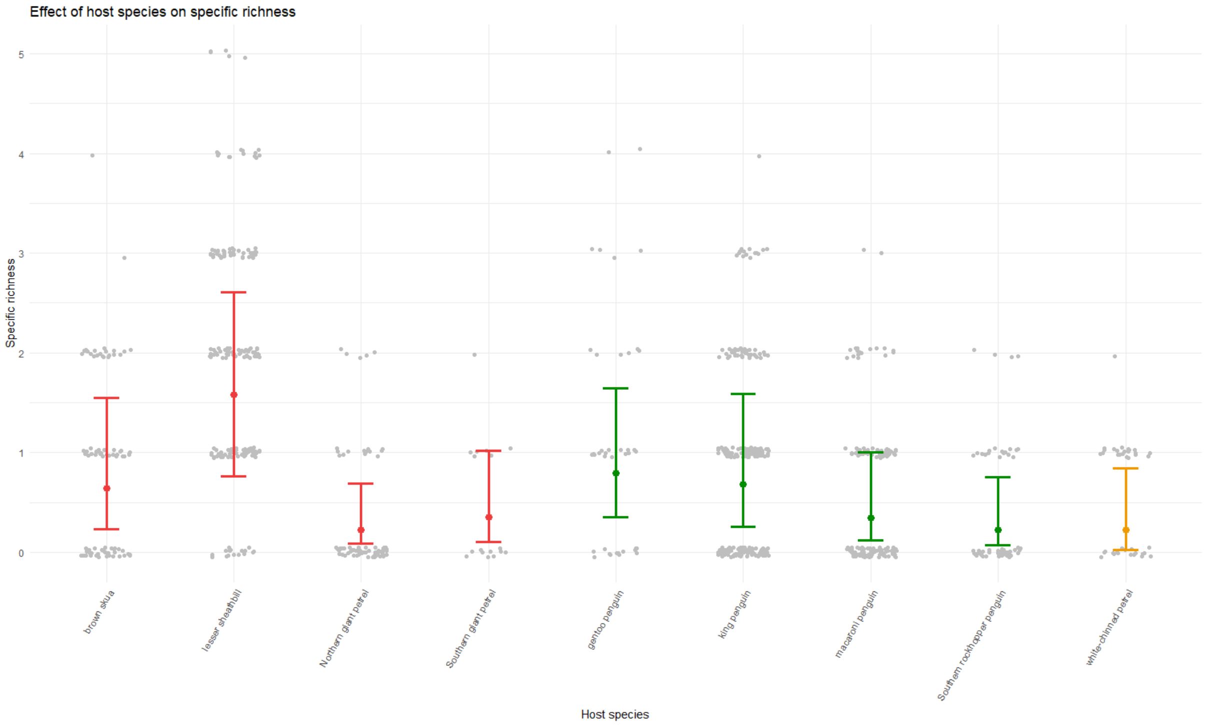

**Supplementary Material – S8| Second-step characterization of *Mycobacterium*-positive samples from this study.** Forty-four samples were screened for environmental infectious agents, mainly *Mycobacterium spp*. and *amoeba*, using a multi-PCR tool (Michelet et al. *unpublished*).

| Sample_ID | Island | Year | Species | *M. bovis* | *M. microti* | *M. kansasii* | *M. intermedium* | *M. bourgelatii* | MAC | *Maa* | *Map* | *M. arupense* | *M. engbaekii* | *M. nonchromogenicum* | *M. terrae* cpx | *M. porcinum* | *M. smegmatis* | *M. moracense* | *M. thermoresistibile* | *M. vaccae* | *Mycobacterium* spp. | *Acanthamoeba* sp. | *Naegleria fowleri* | *Balamuthia mandrillaris* | *Hartmannella vermiformis* | *E.canis* |
|---|---|---|---|---|---|---|---|---|---|---|---|---|---|---|---|---|---|---|---|---|---|---|---|---|---|---|
| AMS22-YNA-048 | Amsterdam | 2022 | yellow-nosed albatross | - | - | 31,64 | - | - | - | - | - | - | - | - | 19,98 | - | - | - | - | - | 17,3 | - | - | - | - | 29,48 |
| AMS22-YNA-048 | Amsterdam | 2022 | yellow-nosed albatross | - | - | 37,66 | - | - | - | - | - | - | - | - | 19,54 | - | - | - | - | - | 17,73 | - | - | - | - | 26,31 |
| AMS22-YNA-075 | Amsterdam | 2022 | yellow-nosed albatross | - | - | - | - | - | - | - | - | - | - | - | 19,03 | - | - | - | - | - | 17,54 | 27,16 | - | - | - | 25,58 |
| AMS22-YNA-075 | Amsterdam | 2022 | yellow-nosed albatross | - | - | - | - | - | - | - | - | - | - | - | 18,39 | - | - | - | - | - | 17,67 | 26,99 | - | - | - | 25,38 |
| AMS22-YNA-083 | Amsterdam | 2022 | yellow-nosed albatross | - | - | - | - | - | - | - | - | - | - | - | 18,85 | - | - | - | - | - | 17,36 | 25,66 | - | - | - | 26,4 |
| AMS22-YNA-083 | Amsterdam | 2022 | yellow-nosed albatross | - | - | - | - | - | - | - | - | - | - | - | 19,08 | - | - | - | - | - | 17,09 | - | - | - | - | 24,44 |
| AMS22-YNA-126 | Amsterdam | 2022 | yellow-nosed albatross | - | - | - | - | - | - | - | - | - | - | - | 19,66 | - | - | - | - | - | 17,93 | - | - | - | - | 24,56 |
| AMS22-YNA-126 | Amsterdam | 2022 | yellow-nosed albatross | - | - | - | - | - | - | - | - | - | - | - | 19,72 | - | - | - | - | - | 17,96 | - | - | - | - | 27,38 |
| AMS22-YNA-159 | Amsterdam | 2022 | yellow-nosed albatross | - | - | 34,71 | - | - | - | - | - | - | - | - | 18,58 | - | - | - | - | - | 14,65 | - | - | - | - | 23,94 |
| AMS22-YNA-159 | Amsterdam | 2022 | yellow-nosed albatross | - | - | 33,08 | - | - | - | - | - | - | - | - | 18,63 | - | - | - | - | - | 14,64 | - | - | - | - | 30,65 |
| AMS22-YNA-162 | Amsterdam | 2022 | yellow-nosed albatross | - | - | - | - | - | - | - | - | - | - | - | 18,95 | - | - | - | - | - | 15,83 | - | - | - | - | 24,07 |
| AMS22-YNA-162 | Amsterdam | 2022 | yellow-nosed albatross | - | - | - | - | - | - | - | - | - | - | - | 19,02 | - | - | - | - | - | 15,91 | - | - | - | - | 26,51 |
| AMS22-YNA-178 | Amsterdam | 2022 | yellow-nosed albatross | - | - | - | - | - | - | - | - | - | - | - | 20,11 | - | - | - | - | - | 17,87 | 25,48 | - | - | - | 26,21 |
| AMS22-YNA-178 | Amsterdam | 2022 | yellow-nosed albatross | - | - | - | - | - | - | - | - | - | - | - | 19,9 | - | - | - | - | - | 17,91 | - | - | - | - | 24,79 |
| AMS22-YNA-206 | Amsterdam | 2022 | yellow-nosed albatross | - | - | - | - | - | - | - | - | - | - | - | 21,3 | - | - | - | - | - | 18,87 | - | - | - | - | 24,92 |
| AMS22-YNA-206 | Amsterdam | 2022 | yellow-nosed albatross | - | - | - | - | - | - | - | - | - | - | - | 20,8 | - | - | - | - | - | 18,63 | - | - | - | - | 26,4 |
| AMS22-YNA-207 | Amsterdam | 2022 | yellow-nosed albatross | - | - | - | - | - | - | - | - | - | - | - | 21,14 | - | - | - | - | - | 18,06 | - | - | - | - | 26,52 |
| AMS22-YNA-207 | Amsterdam | 2022 | yellow-nosed albatross | - | - | - | - | - | - | - | - | - | - | - | 20,92 | - | - | - | - | - | 18,08 | - | - | - | - | 24,18 |
| AMS22-YNA-218 | Amsterdam | 2022 | yellow-nosed albatross | - | - | - | - | - | - | - | - | - | - | - | 21,22 | - | - | - | - | - | 18,9 | - | - | - | - | 27,01 |
| AMS22-YNA-218 | Amsterdam | 2022 | yellow-nosed albatross | - | - | - | - | - | - | - | - | - | - | - | 21,18 | - | - | - | - | - | 18,96 | - | - | - | - | 25,13 |
| AMS22-YNA-276 | Amsterdam | 2022 | yellow-nosed albatross | - | - | - | - | - | - | - | - | - | - | - | 18,06 | - | - | - | - | - | 14,38 | 27,77 | - | - | - | 30,42 |
| AMS22-YNA-276 | Amsterdam | 2022 | yellow-nosed albatross | - | - | 32,75 | - | - | - | - | - | - | - | - | 17,98 | - | - | - | - | - | 14,33 | 26,97 | - | - | - | 26,03 |
| AMS22-YNA-297 | Amsterdam | 2022 | yellow-nosed albatross | - | - | - | - | - | - | - | - | - | - | - | 20,05 | - | - | - | - | - | 17,18 | - | - | - | - | 24,56 |
| AMS22-YNA-297 | Amsterdam | 2022 | yellow-nosed albatross | - | - | - | - | - | - | - | - | - | - | - | 20,19 | - | - | - | - | - | 17,13 | - | - | - | - | 24,44 |
| AMS22-YNA-319 | Amsterdam | 2022 | yellow-nosed albatross | - | - | 32,84 | - | - | - | - | - | - | - | - | 20,26 | - | - | - | - | - | 17,8 | 27,18 | - | - | - | 25,44 |
| AMS22-YNA-319 | Amsterdam | 2022 | yellow-nosed albatross | - | - | 32,73 | - | - | - | - | - | - | - | - | 19,79 | - | - | - | - | - | 17,75 | 27,1 | - | - | - | 30,42 |
| AMS22-YNA-352 | Amsterdam | 2022 | yellow-nosed albatross | - | - | - | - | - | - | - | - | - | - | - | 19,34 | - | - | - | - | - | 17,79 | - | - | - | - | 26,92 |
| AMS22-YNA-352 | Amsterdam | 2022 | yellow-nosed albatross | - | - | - | - | - | - | - | - | - | - | - | 19,35 | - | - | - | - | - | 17,9 | - | - | - | - | 26,24 |
| AMS22-YNA-353 | Amsterdam | 2022 | yellow-nosed albatross | - | - | - | - | - | - | - | - | - | - | - | 20,69 | - | - | - | - | - | 18,61 | - | - | - | - | 25,57 |
| AMS22-YNA-353 | Amsterdam | 2022 | yellow-nosed albatross | - | - | - | - | - | - | - | - | - | - | - | 20,44 | - | - | - | - | - | 18,7 | - | - | - | - | 29,03 |
| AMS22-YNA-408 | Amsterdam | 2022 | yellow-nosed albatross | - | - | - | - | - | - | - | 27,49 | - | - | - | 18,77 | - | - | - | - | - | 16,32 | - | - | - | - | 24,74 |
| AMS22-YNA-408 | Amsterdam | 2022 | yellow-nosed albatross | - | - | - | - | - | - | - | - | - | - | - | 18,66 | - | - | - | - | - | 16,18 | - | - | - | - | 25,37 |
| AMS22-YNA-553 | Amsterdam | 2022 | yellow-nosed albatross | - | - | 32,53 | - | - | - | - | - | - | - | - | 18,33 | - | - | - | - | - | 16,69 | - | - | - | 25,29 | 24,25 |
| AMS22-YNA-553 | Amsterdam | 2022 | yellow-nosed albatross | - | - | 31,5 | - | - | - | - | - | - | - | - | 18,42 | - | - | - | - | - | 16,72 | - | - | - | 26,69 | 25,51 |
| AMS22-YNA-554 | Amsterdam | 2022 | yellow-nosed albatross | - | - | - | - | - | - | - | - | - | - | - | 20,88 | - | - | - | - | - | 19,75 | - | - | - | 26,81 | 27,4 |
| AMS22-YNA-554 | Amsterdam | 2022 | yellow-nosed albatross | - | - | - | - | - | - | - | - | - | - | - | 20,66 | - | - | - | - | - | 19,32 | - | - | - | 26,54 | 24,27 |
| AMS22-YNA-569 | Amsterdam | 2022 | yellow-nosed albatross | - | - | - | - | - | - | - | - | - | - | - | 20,84 | - | - | - | - | - | 18,82 | - | - | - | - | 26,13 |
| AMS22-YNA-569 | Amsterdam | 2022 | yellow-nosed albatross | - | - | - | - | - | - | - | - | - | - | - | 20,88 | - | - | - | - | - | 18,85 | - | - | - | 28 | 24,94 |
| AMS22-YNA-570 | Amsterdam | 2022 | yellow-nosed albatross | - | - | - | - | - | - | - | - | - | - | - | 19,43 | - | - | - | - | - | 17,94 | - | - | - | 25,87 | 24,73 |
| AMS22-YNA-570 | Amsterdam | 2022 | yellow-nosed albatross | - | - | - | - | - | - | - | - | - | - | - | 19,48 | - | - | - | - | - | 17,86 | - | - | - | 26,23 | 24,72 |
| AMS22-YNA-572 | Amsterdam | 2022 | yellow-nosed albatross | - | - | - | - | - | - | - | - | - | - | - | 19,54 | - | - | - | - | - | 18,76 | - | - | - | - | 25,12 |
| AMS22-YNA-572 | Amsterdam | 2022 | yellow-nosed albatross | - | - | - | - | - | - | - | - | - | - | - | 19,39 | - | - | - | - | - | 18,63 | - | - | - | - | 24,97 |
| KER18-BLP-002 | Kerguelen | 2018 | blue petrel | - | - | - | - | - | - | - | - | - | - | - | - | - | - | - | - | - | 22,23 | - | - | - | - | 27,29 |
| KER18-BLP-002 | Kerguelen | 2018 | blue petrel | - | - | - | - | - | - | - | - | - | - | - | - | - | - | - | - | - | 21,93 | - | - | - | - | 30,14 |
